# Dynamic structural changes and inhibition of insect delta and epsilon glutathione *S*-transferases by ethacrynic acid and permethrin

**DOI:** 10.64898/2026.07.24.740637

**Authors:** Meha Sharma, Sibei Qin, Christopher J. Thibodeaux, Mehran Dastmalchi

## Abstract

Insect glutathione *S*-transferases (GSTs) play critical roles in xenobiotic detoxification and insecticide resistance, making them promising targets for selective pest-control strategies. Here, we performed a comparative analysis of GSTs representing multiple classes from beneficial insects, agricultural pests, and disease vectors. We found significant isozyme-specific variations in catalytic activity, stability, and conformational dynamics, notably, in relation to inhibition by the commercial chemical agents, ethacrynic acid (ECA) and permethrin (PER). Sequence similarity network analysis revealed distinct clustering of major GST classes and further highlighted the relatively recent evolutionary divergence of the insect-specific delta and epsilon classes. Structural modeling revealed highly conserved glutathione-binding sites (G-site), but substantial variation in the hydrophobic substrate-binding regions (H-site). Further, epsilon-class GSTs exhibited α4 helices oriented approximately 10° closer to the glutathione-binding site than delta enzymes, suggesting differences in active-site architecture. Steady-state kinetic analyses using 1-chloro-2,4-dinitrobenzene (CDNB) demonstrated that epsilon GSTs are generally more catalytically efficient. Inhibition studies revealed that ECA acts as a potent mixed-type inhibitor of delta-class GSTs, reducing catalytic efficiency by up to 56-fold, whereas PER produced weaker and more species-dependent effects. Notably, ECA binding strongly stabilized delta GSTs, as measured by differential scanning fluorimetry (DSF) and induced a selective rigidification of the α3/α4 linker region and active-site motifs, as observed by hydrogen-deuterium exchange mass spectrometry (HDX-MS). Collectively, these findings demonstrate the importance of analyzing the conformational dynamics mediating insect GST inhibition and provide a framework for exploiting isozyme-specific structural features in the design of next-generation selective insecticides.

## INTRODUCTION

Xenobiotic resistance in insects is rendered through the metabolic capacity of its detoxification systems or reduced target site sensitivity. Three classes of enzymes are responsible for the majority of detoxification: cytochrome P450 monooxygenases (P450s), esterases, and glutathione *S*-transferases (GSTs)^1–5^. Evolution of insecticide resistance is often associated with adaptive genomic changes in the function or expression, or both, of genes encoding this machinery^3^. These enzymes conjugate, neutralize, or metabolize xenobiotics, including plant allelochemicals and synthetic insecticides, facilitating their export, sequestration, or catabolism^6,7^.

Cytosolic GSTs mediate resistance to a range of xenobiotics, including insecticides incorporating organophosphate (OP), organochlorines (OC), and pyrethroids. These enzymes function by conjugating reduced glutathione (GSH) to the electrophilic centers of target compounds ranging from insecticides (e.g., carbonyls, epoxides) to endogenously activated compounds (e.g., lipid peroxidation products). The latter role is crucial for maintaining cellular antioxidant defence against oxidative stress^2,8,9^. There are also examples of GSH acting as a cofactor rather than a substrate for conjugation^10^. GSTs can additionally act as binding proteins, facilitating intracellular and vascular transport or sequestration, particularly of endogenous lipophilic compounds^11–13^. GST gene duplication, functional divergence, and expression changes have been correlated to insecticide resistance. In one example, an organophosphate resistance strain of *Musca domestica* (housefly) carried 12 copies of the *MdGSTd3*-like loci, compared to only 3 in the susceptible strain^14–16^.

Insect GSTs are classified according to amino acid sequence homology, and are assigned Greek letters (corresponding to mammalian GST classes^17^), including delta (formerly class I), epsilon (class III), and sigma (class II)^18^. The delta and epsilon GSTs are often the most numerous classes in insects, with the numbers of isozymes varying between species, mainly due to lineage-specific duplication events^3^. These two classes of GSTs are the most highly implicated in insecticide resistance. Interestingly, the epsilon-class GSTs are enriched in the dipterans, coleopterans, and lepidopterans and do not occur in the hymenoptera (e.g., bees) and exopterygota (e.g., cockroaches)^19^. The absence of the epsilon GSTs has been speculated to account for sensitivity to certain insecticides, e.g., in the crucial pollinator species, *Apis mellifera* (honeybee)^20^. Notably, delta and epsilon GSTs are not found in mammals, further positioning this class of enzymes as a target for novel insecticides^21^. Therefore, the probing of differences in structure, ligand-binding, and catalytic properties between the delta and epsilon GST classes is of crucial importance.

Xenobiotic substrates for insect delta and epsilon GSTs vary depending on the isozyme, but include OP insecticides (e.g., malathion, parathion, and acephate)^11^, OC insecticides (e.g., DDT (dichlorodiphenyltrichloroethane), lindane or γ-hexachlorocyclohexane, dieldrin, endosulfan)^12,14^, pyrethroids (e.g., cyhalothrin, β-cypermethrin)^13,22^, and compounds belonging to other classes (e.g., fipronil, indoxacarb, chlorantraniliprole, spinosad)^6^. The detoxification of OPs involves GSH conjugation through *O*-dealkylation or *O*-dearylation reactions, yielding less toxic derivatives such as *O*-alkyl, *O*-aryl, and phosphonate conjugates^23^. For organochlorine insecticides, detoxification typically involves dehydrochlorination and frequently GSH conjugation of the parent compounds^9^. In contrast, pyrethroids induce oxidative damage in cells, which GSTs help mitigate by detoxifying the resulting lipid peroxidation products^22^.

Cytosolic GSTs are dimeric, with each monomer containing a highly-conserved glutathione-binding site (G-site) and a more structurally varied binding site for hydrophobic toxic compounds (H-site)^23,24^. The *N*-terminal domain, consisting of approximately 80 residues, adopts a thioredoxin-like fold and harbors the G-site^23^. The *C*-terminal domain is largely alpha-helical and contains the highly hydrophobic H-site. Variation in *C*-domain topology is important for substrate selectivity, allowing GST paralogs within the same species to discriminate between substrates^25,26^. Notably, the dimer interfaces of GSTs are also a locus of structural variation, with the delta and epsilon-classes described as lock-and-key ‘clasp’^27^ or ‘wafer’^28^ motifs, respectively. In the delta GSTs, aromatic amino acid side chains from both subunits undergo a pi-stacking interaction to form the ‘key’ that locks the dimer together. Whereas the epsilon ‘wafer’ motif also resides in the active site, with a histidine-serine in the subunit interface, contributing both to quaternary stability as well as directing the two GSH molecules in the opposing active sites^28^.

Another notable characteristic of cytosolic GSTs is their catalytic redundancy, as many of these enzymes can catalyze GSH conjugation with similar electrophilic substrates. Using the model substrate 1-chloro-2,4-dinitrobenzene (CDNB), several studies have suggested that elevated CDNB activity of GSTs correlates positively with resistance via the metabolism or sequestration of xenobiotics^29^. Conversely, certain compounds have been advanced as inhibitors of GST activity, including permethrin (PER) and ethacrynic acid (ECA); however, their mode of binding and inhibition has been difficult to ascertain.

PER is a pyrethroid-type insecticide that functions through neuronal disruption and incapacitates insects before they can rupture the skin. Inhibition studies of susceptible insect GSTs by PER have yielded contradictory outcomes. In *Anopheles* spp. (mosquito), competitive, non-competitive, and uncompetitive inhibition have all been reported, depending on the enzyme variant^8,30^. In one striking example, PER inhibition of CDNB activity of three splice variants of *Anopheles dirus* delta-class GSTs were characterized as uncompetitive (AdGST1-2), non-competitive (AdGST1-3), and competitive (AdGST1-4)^31^. These distinct outcomes suggest that subtle structural differences in GSTs can result in varying modes and magnitude of inhibition by insecticides.

The phenoxyacetic acid derivative ECA (trade name Edecrin) is a potent reversible inhibitor of human and rat GSTs^32^. Notably, the inhibitory effect of ECA on human GSTs is speculated to sensitize human cancer cells to the effects of anticancer agents^33^. Commercially, it is widely used as a diuretic drug to treat high blood pressure, inflammation, and swelling. The mode of action is not entirely understood; however, ECA contains an α,β-unsaturated ketone, which has been implicated in GST-mediated inhibition^34^. It has also been suggested that ECA can reduce GSH levels through adduct formation, limiting GSH availability to GSTs. Despite these investigations, the exact inhibitory mechanisms and roles of ECA are poorly understood and could vary in different contexts.

In this work, we conducted an in-depth biochemical and biophysical analysis of selected insect GST enzymes to probe differences in structure, enzyme kinetics, and inhibition properties. Our analysis spanned three classes of GSTs (delta, epsilon, and sigma) from insect pests, disease vectors, and beneficial species. Our analysis focused on *Apis mellifera* (Ap) and *Bombyx mori* (Bm), which are also commercially beneficial as primary producers of honey and silk, respectively; agricultural pests, including Colorado potato beetle, *Leptinotarsa decemlineata* (Ld), a major pest of potato, and diamondback moth *Plutella xylostella* (Px), which feeds on the vegetation of the Brassica family (e.g., cauliflower, cabbage, and broccoli). Finally, we have included a GST from *Aedes aegypti* (Aa), which is a vector for diseases, including dengue fever, yellow fever, and chikungunya. Using a combination of assays investigating steady state kinetics and inhibition of GSTs, we showed that PER and ECA exhibit distinct isozyme-and species-specific inhibition properties. We further explored the biophysical basis of inhibition with differential scanning fluorimetry (DSF) and hydrogen-deuterium exchange mass spectrometry (HDX-MS). Cumulatively, the results highlight the varied and dynamic nature of GST function and inhibition and suggest the potential for novel insecticides tailored to GST isozymes.

## RESULTS

### Bioinformatic analysis of insect cytosolic GST enzymes

Our target enzymes include delta isozymes from *B. mori* (BmGSTd2), *P. xylostella* (PxGSTd3), *L. decemlineata* (LdGSTd3), and *A. mellifera* (ApGSTd1), epsilon class enzymes from *A. aegypti* (AaGSTe2), *B. mori* (BmGSTe3), and *P. xylostella* (PxGSTe4), and a sigma class enzyme from *B. mori* (BmGSTs1). To better visualize the sequence relationships between our target enzymes and the broader GST superfamily, we created a sequence similarity network of all insect GSTs using the Enzyme Function Initiative Enzyme Similarity Tool (EFI-EST)^35^. The initial network was generated for the entire cytosolic GST family (Interpro ID = IPR040079) using the UniRef90 database and a phylogenetic filter for class Insecta. In the UniRef90 database, all sequences with >90% identity were grouped into a single entry. This step was necessary due to the large size of the IPR040079 family (186,670 at the time of this study). Once the initial set of 3,003 insect GST UniRef90 entries were retrieved, the Cluster Analysis tools of the EFI were used to expand the UniRef90 entries into individual UniProt IDs (a total of 4,725 unique sequences). These 4,725 UniProt sequences were then used to generate the final network at an alignment score (*E*-value) of 65 (**Fig. 1A**).

**Fig. 1.**
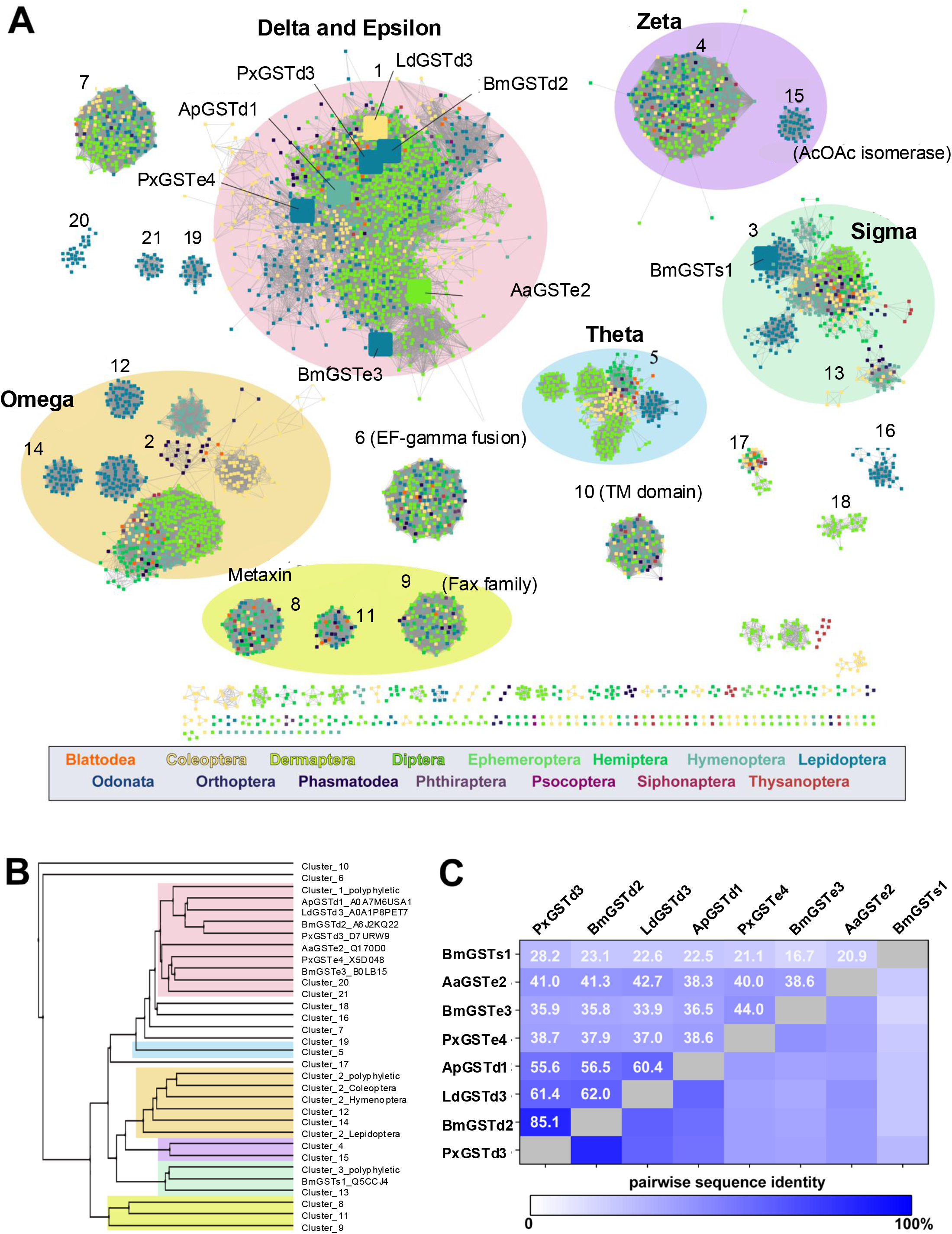
Phylogenetic analysis of insect GSTs. **A.** Sequence similarity network of all GSTs from class Insecta. Nodes (individual enzymes) are coloured according to phylogenetic order. The delta and epsilon enzymes examined in this study are labeled. **B.** Phylogenetic tree of the major sequence clusters (numbered 1-21 in panel A). **C.** Pairwise amino acid sequence identity of the GSTs investigated in this study showing the greater shared identity among the delta isozymes.

The curated insect GST proteins (coloured according to taxonomic order in **Fig. 1A**) clustered into distinct clades representing all major GST structural families known to exist in class Insecta including the delta, epsilon, sigma, omega, theta, and zeta classes. Other major clusters in the GST family include failed axon connection (FAX) homologues, metaxin (MTX) like proteins involved in mitochondrial membrane protein transport and insertion, and the microtubule binding protein EF1-γ. To better observe the relationships between the major GST classes, we generated and aligned consensus sequences (**Fig. S1-2**) for each of the major GST clusters containing more than 20 nodes (**Fig. 1B**). Some of the GST classes cluster into tight groups of mixed phylogenies such as the zeta, FAX, and metaxin groups, suggesting a high level of similarity within these clusters across all insects. In contrast, the sigma, theta, and omega groups parse largely according to phylogeny, in which individual orders form distinct subclusters within the group. This hints at the possibility of structural diversification among GSTs that may enable order-specific insecticide development. The insect-specific delta and epsilon enzymes form a single large cluster, consistent with their more recent evolutionary divergence^19^. At higher alignment scores (shown in **Fig. S1**), the delta enzymes can be parsed from the epsilon enzymes suggesting that the delta enzymes share more sequence identity than the epsilon enzymes. The target GST enzymes in this study are highlighted in the network and their pairwise sequence identities are provided in **Fig. 1C**.

### Structural modeling of selected GSTs

High resolution structures are available for several insect delta and epsilon GSTs (**Fig. S2**), including BmGSTd2 (PDB: 3VK9) among our target enzymes^36,37^. Alignment of amino acid sequences from our selected GSTs (**Fig. S3**) confirms a high degree of conservation in the G-site, including the NPQHTVPTL and D/E-S-H/R motifs involved in GSH binding, whereas the H-site residues exhibit greater variation. These enzymes are all predicted to form homodimers with each monomer characterized by an *N*-terminal thioredoxin-like domain and a *C*-terminal α-helical domain. Using our sequence similarity network of all insect delta/epsilon enzymes, we performed structural alignments of AlphaFold 3 (AF3) models of our target enzymes with their nearest network neighbors of known structure. Excellent alignments were obtained between our target enzymes and enzymes of known structure (RMSDs ranging from 0.233-1.254 Å) (**Fig. S4**; **Table S1**), suggesting that the AF3 models should allow reliable interpretation of the kinetic and biophysical data presented below.

To gain a better understanding of ligand binding to our target enzymes, we aligned the AF3 model of BmGSTe3 (**Fig. 2A**) with the epsilon-class GST from *Drosophila melanogaster* (DmGSTe14, PDB ID: 7DB4), which was solved in the presence of both reduced glutathione (GSH) and the hydrophobic substrate analog H2C (**Fig. 2B**). To our knowledge, this is the only high-resolution structure of an insect GST resolved with both G-site and H-site simultaneously occupied with a ligand. In our structural alignment of BmGSTe3 with DmGSTe14, GSH localized within the G-site in close proximity to the active site residues His52, Asp66, Ser67, and His68 (**Fig. 2C**), consistent with the highly conserved G-site structure. In contrast, a steric clash was predicted between H2C and the α4 helix of BmGSTe3, which forms the H-site along with α8 (**Fig. 2D**).

**Fig. 2.**
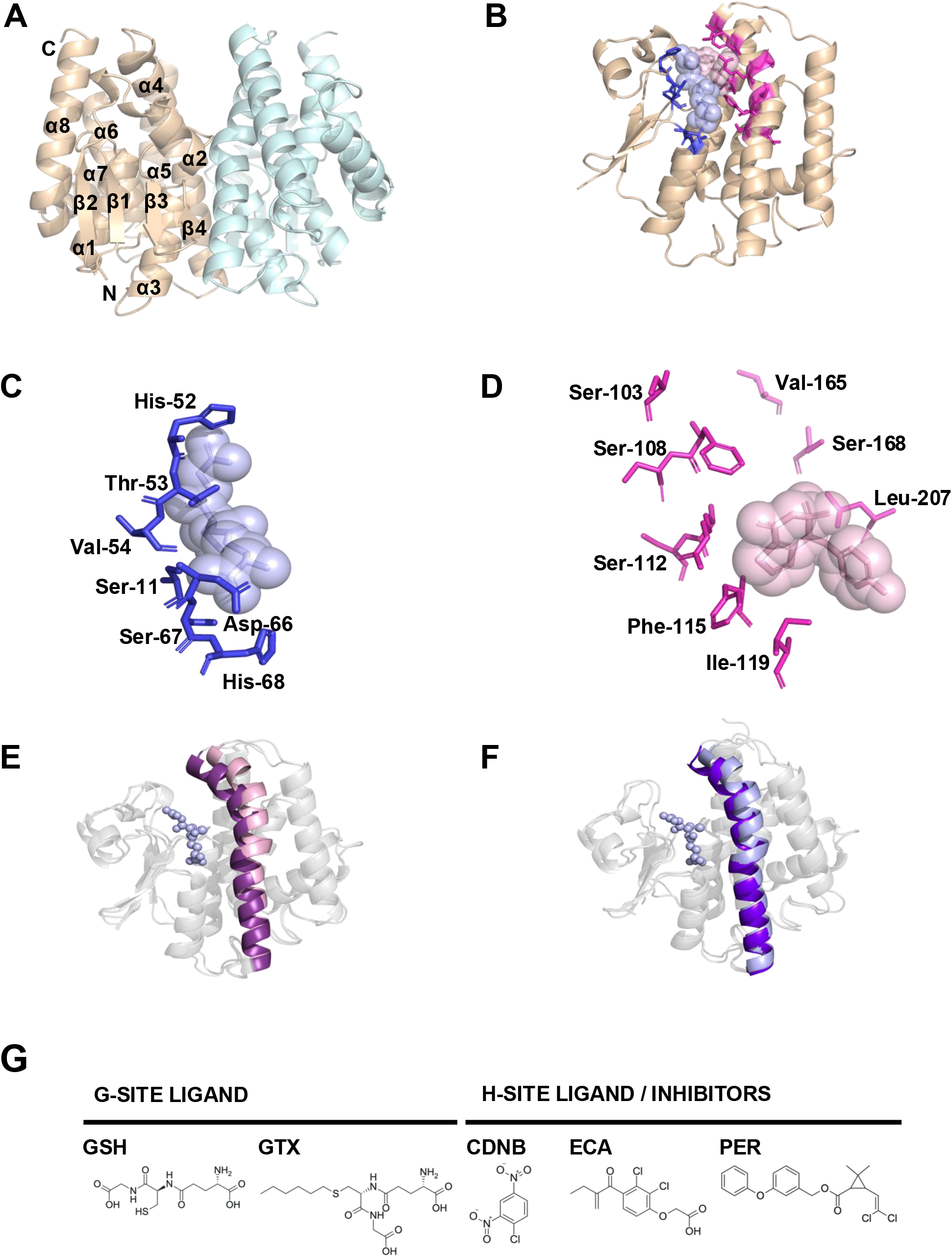
Structural analysis of BmGSTe3. **A.** Predicted dimeric structure of BmGSTe3. The two monomeric units are shown in wheat and pale cyan, respectively. The nomenclature of secondary structural elements is indicated and can also be found in **Fig. S3**. **B.** Structural alignment of BmGSTe3 with the *Drosophila melanogaster* epsilon GST (PDB ID: 7DB4). The background of 7DB4 is hidden for clarity. Ribbon representation of a BmGSTe3 monomer highlighting two bound ligands; glutathione (GSH) shown in light blue and the ligand H2C shown in light pink. Both ligands are depicted using space-filling (sphere) representations to illustrate their occupancy and interaction with active site residues (stick). **C.** Close-up view of GSH binding at the predicted “G-site,” showing detailed interactions within the polar pocket of the enzyme. **D.** Close-up view of H2C binding at the predicted “H-site” (hydrophobic substrate-binding site), indicating its interaction within the hydrophobic channel. Numbers correspond to the residues in BmGSTe3. **E.** Structural alignment of BmGSTd2 and BmGSTe3, featuring their α4-helices, coloured in pink and violet, respectively; GSH ligand is shown in light blue against the grey enzyme. **F.** Structural alignment of PxGSTd3 and PxGSTe4, with α4-helices shown in light blue and purple, respectively. **G.** Chemical structures of ligands used in this study: reduced glutathione (GSH), *S*-hexylglutathione (GTX), 1-chloro-2,4-dinitrobenzene (CDNB), ethacrynic acid (ECA), permethrin (PER).

It has been previously suggested that structural changes in the α4 helix are associated with the functional divergence of delta and epsilon GSTs. The AF3 models of our selected enzymes confirm that epsilon-class enzymes possess α4 helices inclined further towards the substrate (**Fig. 2E**, **F**; **Fig. S5**; **Table S2**). The angles of separation ranged from 41° to 47° for the epsilon enzymes and from 56 to 58° for the delta enzymes (calculations explained in **Materials and Methods**). In our models, other critical residues in the α4 helix (BmGSTe3 numbering: Arg111, Phe115, and Ile119) are brought closer to the GSH-binding site. This is aligned with their previously speculated roles that include forming a hydrogen-bond network that stabilizes the GSH thiolate and rendering a cap over the pocket^38^. Interestingly, in the solved structure of DmGSTe14, the *C*-terminal end of α4 moves away from the G-site to accommodate H2C binding. This movement is accompanied by a re-structuring of the α4-α5 linker (**Fig. S5**). Finally, we noted minor differences in the length of α8-helices, with the epsilon-class enzymes being 1-2 residues longer in *B. mori* and *P. xylostella* than the delta-class. Further, PxGSTe4 had an additional 6 residues at the C-terminus, after the helix.

### Steady state kinetics and inhibition analysis of insect GSTs

We determined the steady state kinetic parameters for our selected insect GSTs (**Fig. S6**) using the substrates 1-chloro-2,4-dinitrobenzene (CDNB) and reduced glutathione (GSH) (**Fig. 2G**; **Fig. 3**; **Table 1**). Reaction of CDNB with GSH yields *S*-2,4-dinitrophenyl glutathione, which is quantified by measuring 340 nm absorbance^39^. *In vitro*, GSTs follow a random sequential bi-bi kinetic mechanism^40–43^, but at physiological GSH concentrations, the mechanism is likely ordered sequential, with GST binding first to the G-site, followed by the electrophilic substrate binding to the H-site. For this reason, we held the GSH concentration in our assays constant at 5 mM and varied the CDNB concentration (50-3200 μM). Our panel of enzymes displayed kinetic parameters for CDNB that are similar to GST enzymes from insects^36,44–47^ and other sources^41,42,48,49^, with K_m,CDNB_ in the 10^-3^-10^-4^ M range, k_Cat_ values in the 10^1^-10^2^ s^-1^ range, and catalytic efficiencies (k_Cat_⁄K_m_) in the range of 10^3^-10^4^M^-1^s^-1^. Our parameters for BmGSTd2 and the Px enzymes agree well with the values reported by Yamamoto and Chen, respectively^36,45^. Overall, the epsilon isozymes in our set had lower *K*_m,CDNB_ values (ranging from 50-225 μM) and higher catalytic efficiencies than the delta enzymes. The BmGSTe3 enzyme had the highest overall catalytic efficiency (430,000 M^-1^s^-1^) in our set, while the BmGSTs1 enzyme had the lowest (3,400 M^-1^s^-1^).

**Fig. 3.**
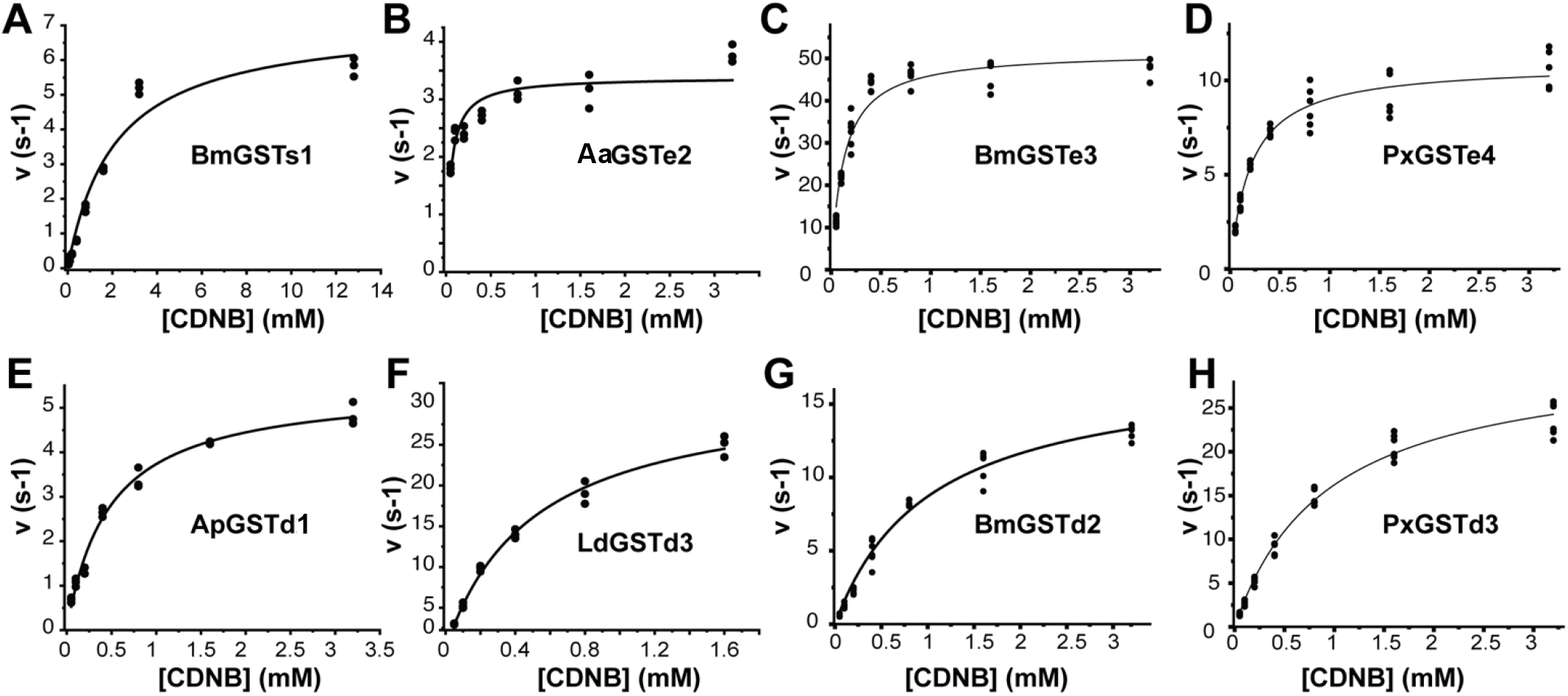
Steady state kinetics of insect GSTs. Enzyme assays were conducted with varying concentrations of CDNB (0.05-3.2 mM), GSH (5 mM), and 35-72 nM of purified recombinant enzyme. ApGSTd1: *Apis mellifera*, BmGSTd2: *Bombyx mori*, LdGSTd3: *Leptinotarsa decemlineata*, PxGSTd3: *Plutella xylostella*, AaGSTe2: *Aedes aegypti*, BmGSTe3, PxGSTe4, BmGSTs1. Data were fitted with the Michaelis-Menten equation. The fitted parameter values are summarized in **Table 1**.

**Table 1:** Steady state kinetic parameters for selected insect glutathione *S*-transferases. Aa: *Aedes aegypti*; Ap: *Apis mellifera*; Bm: *Bombyx mori*; Ld: *Leptinotarsa decemlineata*; Px: *Plutella xylostella*; d: delta; e: epsilon; and s: sigma subclass GST. Values are presented as mean ± standard error.

| Enzyme | $k_{cat}(s^{-1})$ | $K_m(mM)$ | $k_{cat}/K_m(M^{-1}s^{-1})$ |
| --- | --- | --- | --- |
| ApGSTd1 | $5.5 \pm 0.15$ | $0.47 \pm 0.04$ | 11,700 |
| BmGSTd2 | $17.5 \pm 0.56$ | $1.02 \pm 0.08$ | 17,200 |
| LdGSTd3 | $32.0 \pm 1.10$ | $0.50 \pm 0.04$ | 64,000 |
| PxGSTd3 | $31.4 \pm 0.90$ | $0.93 \pm 0.07$ | 33,700 |
| AaGSTe2 | $3.4 \pm 0.10$ | $0.05 \pm 0.01$ | 68,000 |
| BmGSTe3 | $51.6 \pm 1.10$ | $0.12 \pm 0.01$ | 430,000 |
| PxGSTe4 | $10.9 \pm 0.26$ | $0.23 \pm 0.02$ | 47,400 |
| BmGSTs1 | $7.2 \pm 0.36$ | $2.10 \pm 0.28$ | 3,400 |

For detailed analysis of GST inhibition by ECA and PER, we selected delta and epsilon isozymes from *B. mori*, a beneficial insect (BmGSTd2 and BmGSTe3) and from *P. xylostella*, an agricultural pest insect (PxGSTd3 and PxGSTe4). The kinetic data are summarized in **Fig. 4** (ECA inhibition), **Fig. 5** (PER inhibition), and **Table 2**; details of the data analysis are provided in the **Materials and Methods**. In all cases, we observed either non-competitive (reduced k_Cat_) or mixed-type inhibition (reduced k_Cat_ and increased K_m_). This suggests that (depending on the GST) both ECA and PER can bind to either the [E:GSH] or [E:GSH:CDNB] Michaelis complex at sites that are distinct from the CDNB binding site. Importantly, non-competitive and mixed-type inhibitors may allow for greater discrimination between pest and non-pest targets over competitive inhibitors, which would likely broadly target many cellular GSTs. ECA produces a mixed-type inhibition pattern for all enzymes except PxGSTe4 (which showed a non-competitive pattern). It also produced notably stronger inhibition of the delta enzymes (reductions in k_Cat_⁄K_m_of 34-and 56-fold for BmGSTd2 and PxGSTd3, respectively) (**Table 2**) than with the epsilon enzymes (reductions in k_Cat_⁄K_m_of only 5.3-and 3.9-fold for BmGSTe3 and PxGSTe4, respectively). The enhanced inhibition of the delta enzymes stems largely from a significantly increased *K*_m,CDNB_ that was not observed in the epsilon isozymes. In contrast, PER provided an opposite trend, with a non-competitive mode of inhibition on all enzymes except BmGSTe3 (which experienced mixed-type inhibition). PER inhibition was slightly stronger on the Bm enzymes (reductions in k_Cat_⁄K_m_of 4.4-and 4.9-fold for BmGSTd2 and BmGSTe3, respectively) than on the Px enzymes (reductions in k_Cat_⁄K_m_of 1.8-and 2.7-fold for PxGSTd3 and PxGSTe4, respectively). The distinct inhibition patterns induced by ECA and PER suggest that the two inhibitors likely exploit different binding sites in each enzyme.

**Fig. 4.**
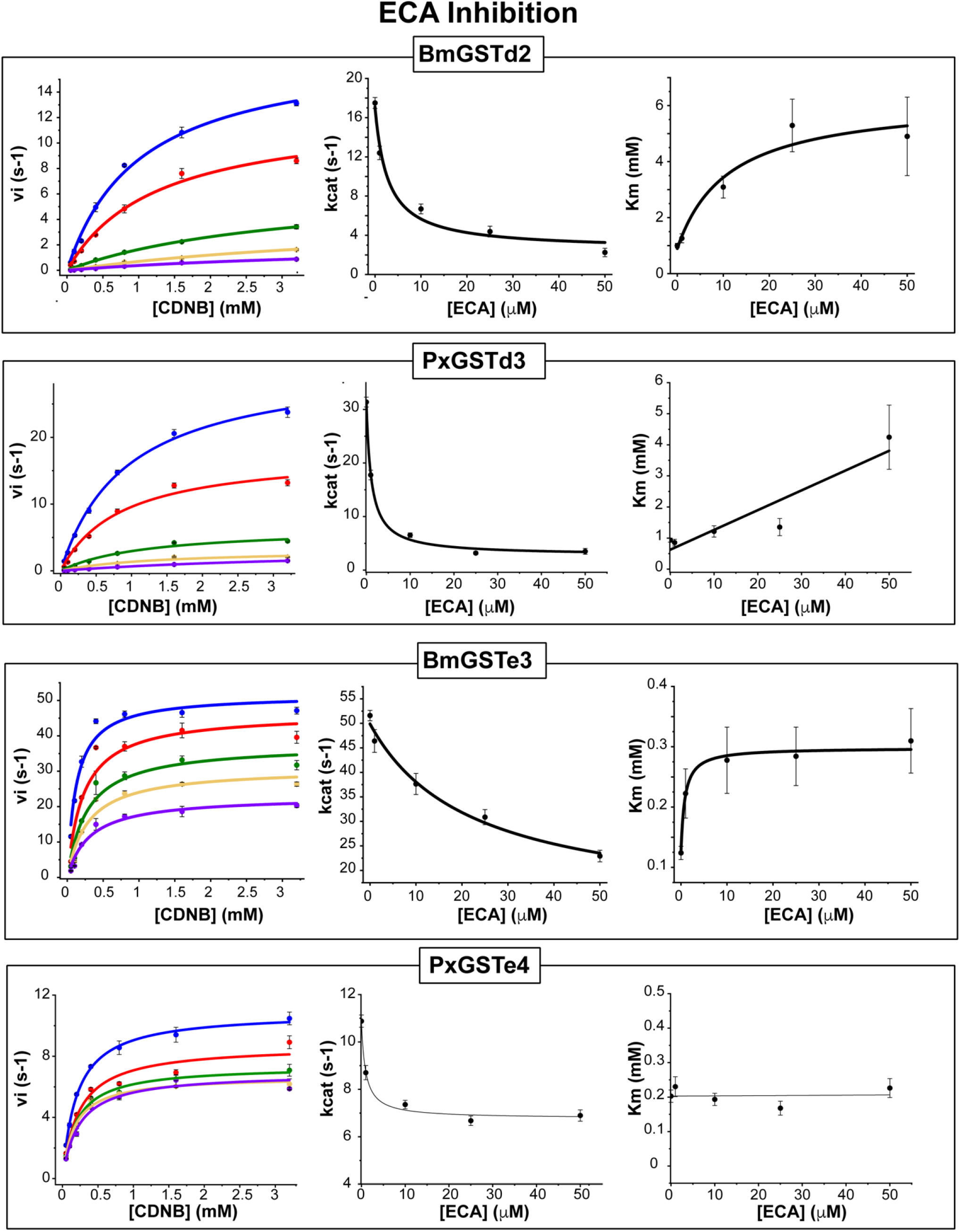
**Inhibition of insect GSTs by ECA**. Steady state kinetic parameters of *Bombyx mori* (Bm) and *Plutella xylostella (*Px) delta and epsilon GSTs for conjugation of GSH-CDNB in the presence of variable concentrations of ECA. From left to right: Michaelis-Menten curves, k_Cat_, and K_m_ plots with increasing inhibitor concentration. [Inhibitor] = 0 (blue), 1 μM (red), 10 μM (green), 25 μM (gold), 50 μM (purple). Outcomes are summarized in **Table 2**.

**Fig. 5.**
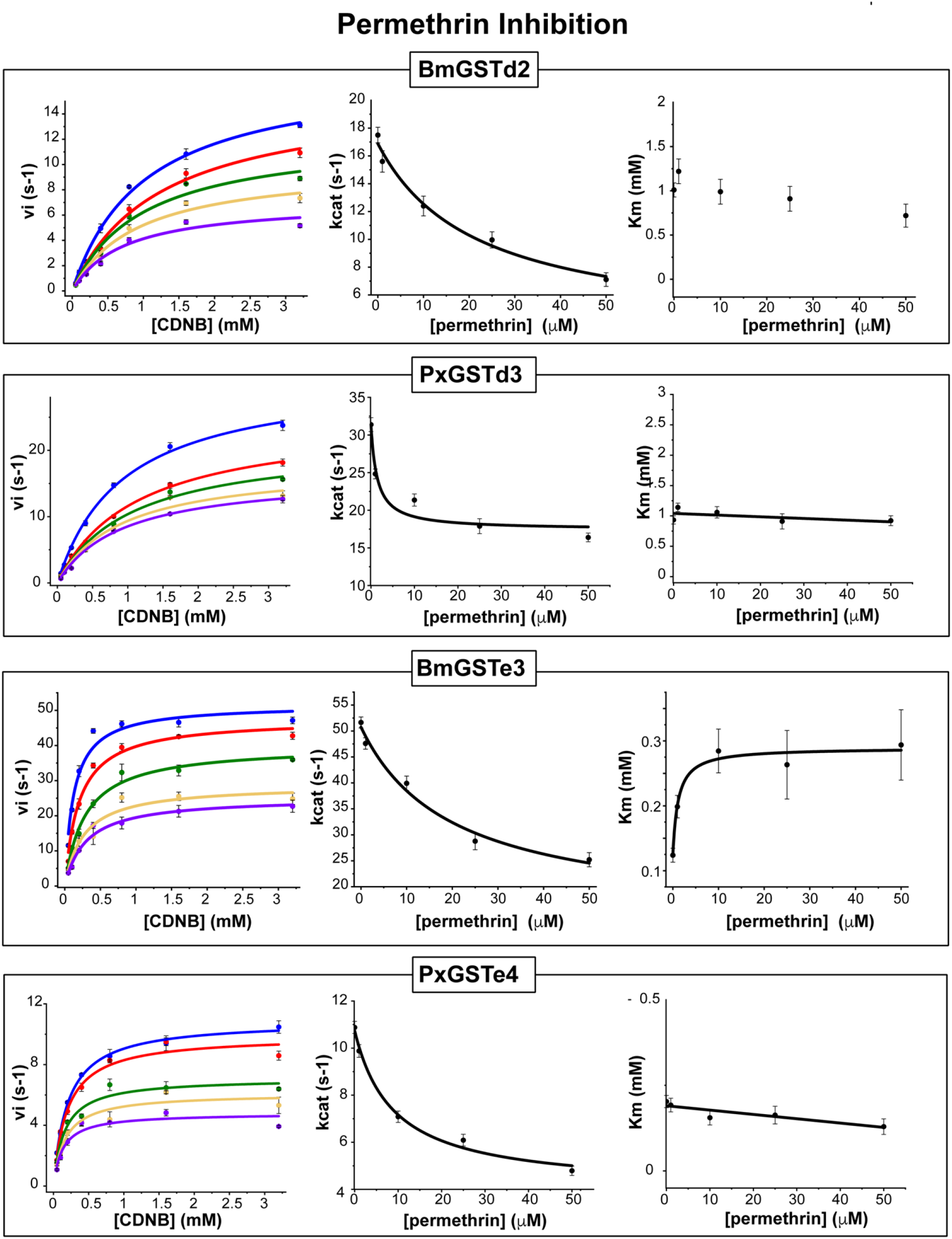
**Inhibition of insect GSTs by PER**. Steady state kinetic parameters of *Bombyx mori* (*Bm*) and *Plutella xylostella* (*Px*) delta and epsilon GSTs for conjugation of GSH-CDNB in the presence of variable concentrations of PER. From left to right: Michaelis-Menten plots, k_Cat_, and K_m_ plots with increasing inhibitor concentration. [Inhibitor] = 0 (blue), 1 μM (red), 10 μM (green), 25 μM (gold), 50 μM (purple). Outcomes are summarized in **Table 2**.

**Table 2:** Ethacrynic acid (ECA) and permethrin (PER) inhibition kinetics of Bm and Px delta and epsilon GSTs.

| Enzyme | Inhibitor | $K_m$ (mM)<br>(no inhibitor) | $K_m$ (mM)<br>(fully inhibited)* | $k_{cat}$ ($s^{-1}$)<br>(no inhibitor) | $k_{cat}(s^{-1})$<br>(fully inhibited)* | Reduction in<br>$k_{cat}/K_m$ | Inhibition<br>Type |
| --- | --- | --- | --- | --- | --- | --- | --- |
| BmGSTd2 | ECA | $1.02 \pm 0.078$ | $5 \pm 1.00$ | $17.5 \pm 0.56$ | $2.5 \pm 0.26$ | 34 | mixed |
| BmGSTe3 | | $0.12 \pm 0.011$ | $0.17 \pm 0.02$ | $52 \pm 1.10$ | $14 \pm 2.70$ | 5.3 | mixed |
| PxGSTd3 | | $0.93 \pm 0.067$ | $4 \pm 1.00^{**}$ | $31.4 \pm 0.90$ | $2.4 \pm 0.08$ | 56 | mixed |
| PxGSTe4 | | $0.23 \pm 0.018$ | no change | $10.9 \pm 0.26$ | $2.8 \pm 0.20$ | 3.9 | noncompetitive |
| BmGSTd2 | PER | $1.02 \pm 0.078$ | no change | $17.5 \pm 0.56$ | $4 \pm 0.80$ | 4.4 | noncompetitive |
| BmGSTe3 | | $0.12 \pm 0.011$ | $0.17 \pm .02$ | $52 \pm 1.10$ | $15 \pm 2.60$ | 4.9 | mixed |
| PxGSTd3 | | $0.93 \pm 0.067$ | no change | $31.4 \pm 0.90$ | $17 \pm 1.50$ | 1.8 | noncompetitive |
| PxGSTe4 | | $0.23 \pm 0.018$ | no change | $10.9 \pm 0.26$ | $4.1 \pm 0.36$ | 2.7 | noncompetitive |

\*determined from fitting the inhibitor induced changes in $k_{cat}$ and $K_m$ to hyperbolic equations as described in the Methods.
\*\*determined at [ECA] = 50 μM

### ECA induces a positive thermal shift in the GST-GSH complex

Differential scanning fluorimetry (DSF) reports on the thermal denaturation and stability of proteins and can be used to assess ligand-induced enzyme structural stabilization. We employed DSF to study the effects of ligand binding on the thermal stability of our Bm and Px delta and epsilon GSTs. Apo enzymes had melting temperatures (T_m_) in the range of 49.5 – 52.0 °C, which increased by 1.0 – 4.5 °C upon GSH binding (**Fig. 6**). The Bm isozymes experienced a greater positive shift than the Px enzymes. The structural basis for this shift is unknown, but the data suggest that the delta enzymes may be slightly more disordered in the absence of GSH ligand. The [E:GSH] complex was subsequently used as a reference point to record the thermal shift (ΔT_m_) induced by the binding of inhibitors (**Fig. 6A, B**).

**Fig. 6.**
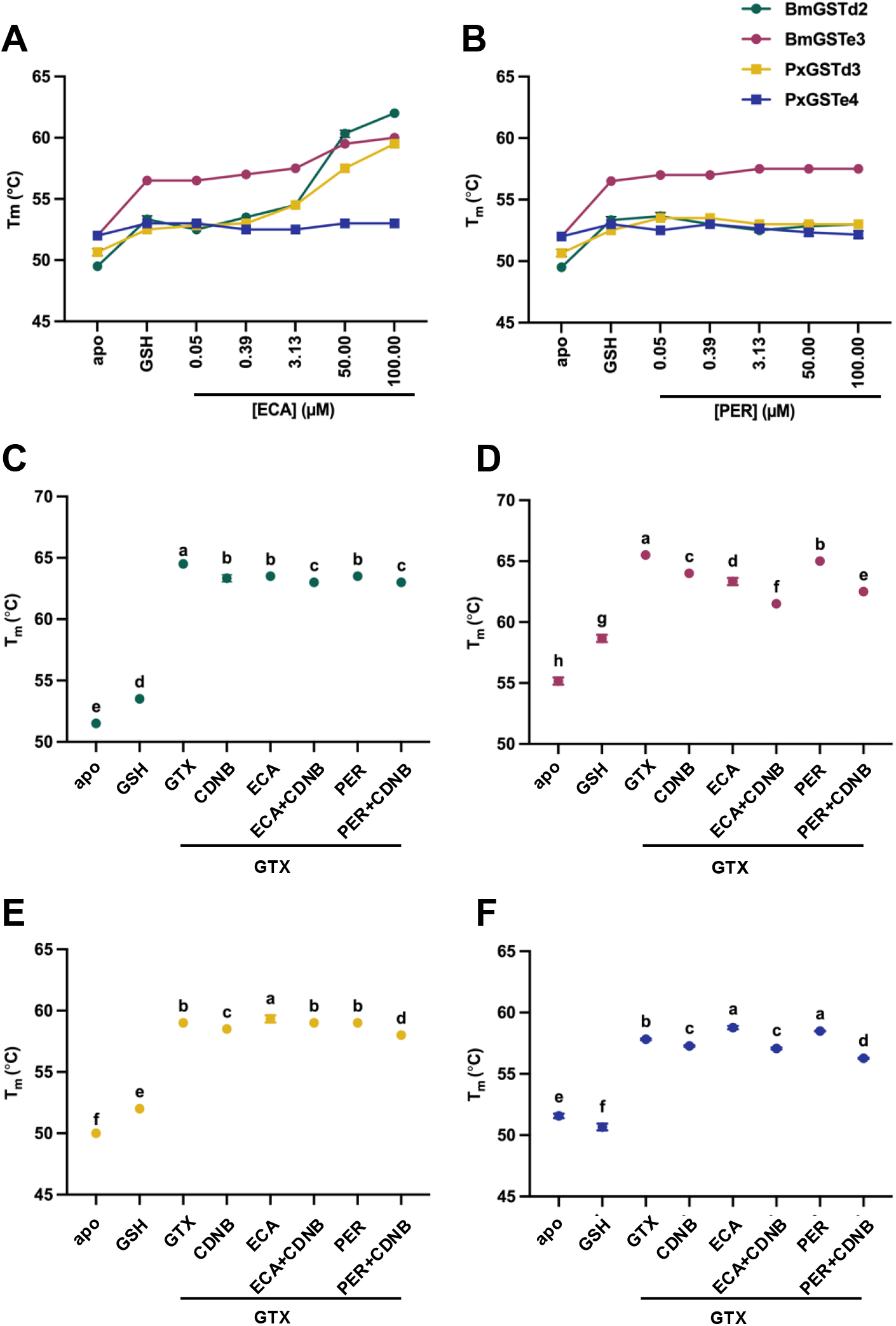
Thermal stability measured by differential scanning fluorimetry (DSF). Increasing concentrations of inhibitors, **A.** ECA and **B.** PER, at a fixed concentration of GSH, measuring T_m_ for each protein mixture. Graphs **C.** BmGSTd2, **D.** BmGSTe3, **E.** PxGSTd3, and **F.** PxGSTe4, represent T_m_ for each protein mixture, under apo (no ligand) condition or with a series of ligands. All values represent the mean ± S.D. (*n*□=□3).

The binding of ECA to the [E:GSH] complex induced significant thermostabilization in all isozymes, except for PxGSTe4. Curiously, PxGSTe4 exhibited the weakest ECA inhibition and was the only enzyme in our set whose K_m_ was insensitive to ECA (**Table 2**). In another interesting parallel with the kinetic inhibition data, the two delta isozymes (BmGSTd2 and PxGSTd3), showed the greatest positive thermal shift upon ECA binding (ΔT_m_ of 8.7 and 7.0 °C, respectively), aligning well with the strong ECA-induced reduction in catalytic efficiency in these enzymes. These data suggest that DSF may provide a powerful high throughput assay for rapid assessment of insect GST susceptibility to ECA inhibition.

Conversely, PER failed to produce a major thermal shift in any of the tested GSTs, with ΔT_m_ values generally within ±1 °C (relative to the [E:GSH]), and no clear concentration dependence (**Fig. 6B**). Again, these DSF data appear to align well with the PER inhibition data. Namely, the insensitivity of K_m,CDNB_ and comparatively weak reduction in catalytic efficiency by PER seem to correspond with the modest ΔT_m_ perturbations. Altogether, the inhibition kinetics and DSF data suggest that ECA and PER bind to insect delta and epsilon GSTs via fundamentally distinct interactions, that trigger distinct effects on enzyme structural stability.

To probe the impact of CDNB binding on thermal stability, we replaced GSH with the inert analogue *S*-hexylglutathione (GTX) to avoid catalysis during the DSF measurement. Structural analysis of several insect GST enzymes has shown that GTX binds to the GSH site^38,50,51^. In contrast to GSH, which produced a modest increase in ΔT_m_ of 1.0 – 4.5 °C, GTX resulted in a drastic increase in thermal stability for all four GSTs (ΔT_m_ of 6.3 – 13.0 °C) (**Fig. 6C-F**). Relative to other ligand additions, GTX established the most significant thermal stabilization. For the Bm enzymes, the addition of ligands (substrate or inhibitor) to the [E:GTX] complex, uniformly resulted in a negative thermal shift (destabilization), particularly when CDNB was included. For the Px enzymes, ligand binding to the [E:GTX] complex produced minimal thermal shifts.

### Detailed HDX-MS analysis of delta and epsilon GSTs

To determine how inhibitor binding modulates the conformational dynamics of our Bm and Px delta and epsilon enzymes on local spatial scales, we employed bottom-up, continuous exchange hydrogen-deuterium exchange mass spectrometry (HDX-MS). Our workflow provided 82-96% sequence coverage depending on the GST with an average redundancy of 3.46 HDX peptides per amino acid residue. Comparative HDX-MS studies were performed by investigating ligand (CDNB, ECA, and PER) binding to the [E:GTX] complexes. Statistically significant differences in deuterium uptake (Welch’s t-test, *p* < 0.01) between ligand bound states were determined as described in the **Materials and Methods**. These changes are mapped onto AF3 models of the target enzymes (**Fig. 7**) and are provided as heatmaps (**Fig. S7**). Coverage maps for all enzymes in all states are provided in **Fig. S8-S11**. Additional raw data and significance calculations are provided in the **Supplementary Information.**

**Fig. 7.**
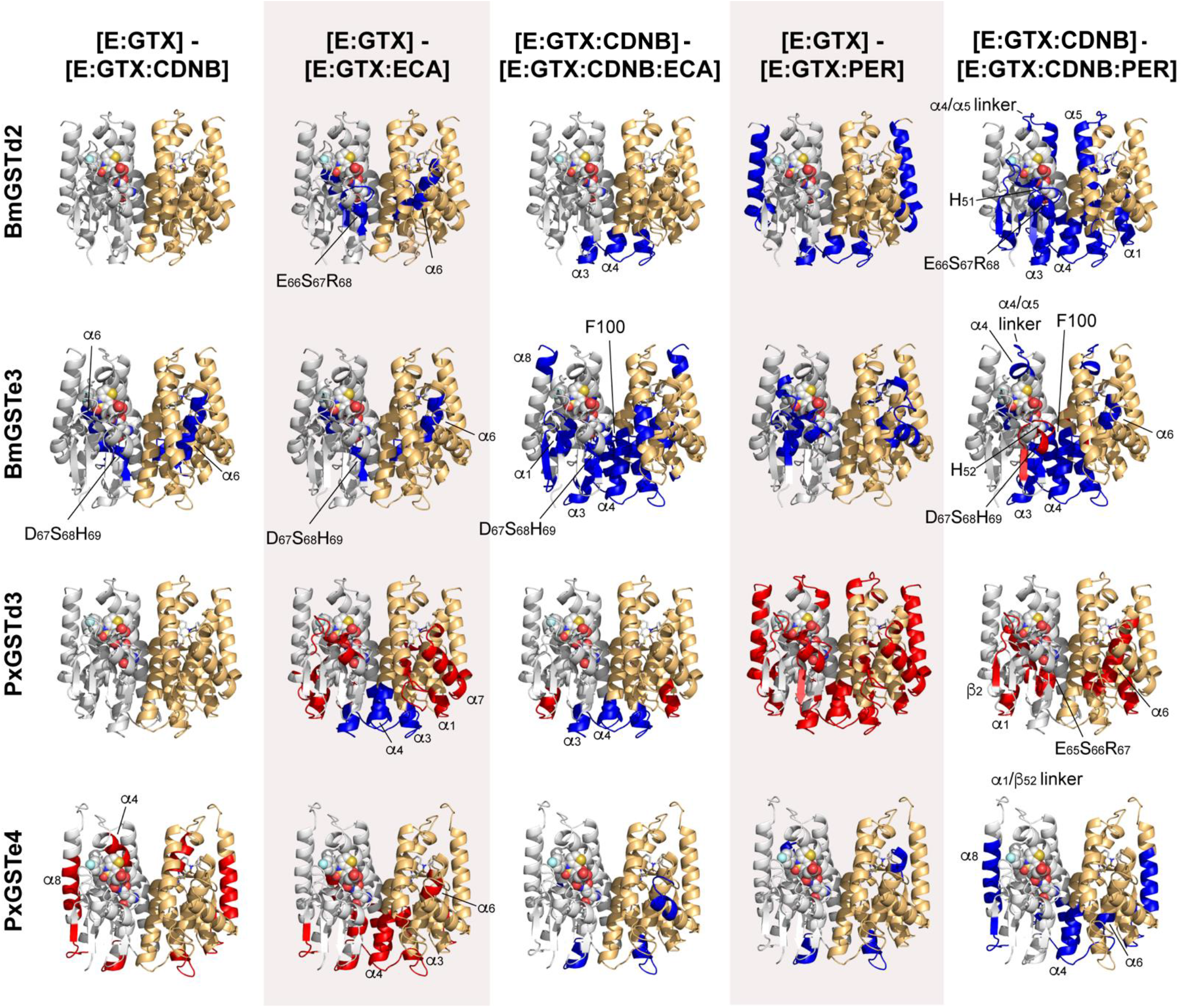
HDX-MS analysis of dynamic conformational changes in delta and epsilon GST enzymes of *B. mori* and *P. xylostella*. Relative deuterium uptake differences were calculated for each enzyme by subtracting the uptake measured for the pairs of biochemical states indicated at the top of the figure. Shown from left to right are the effects caused by CDNB binding to the [E:GTX] complex, ECA binding to the [E:GTX] complex, ECA binding to the [E:GTX:CDNB] complex, PER binding to the [E:GTX] complex, and PER binding to the [E:GTX:CDNB] complex. Significant uptake differences (*p* < 0.01) between states were determined as described in the Materials and Methods and are plotted on AF3 models of the Bm and Px GSTs. Blue/red regions represent decreased/increased deuterium uptake, respectively, according to the difference calculations shown at the top of the figure. These HDX data are shown in heat map format in **Fig. S7** and in coverage map format in **Fig. S8-S11**. HDX changes associated with binding of GTX to the free enzymes are provided in **Fig. S12**.

In agreement with the DSF studies, the binding of GTX to apo-GST resulted in protection from HDX across nearly the entire enzyme structure in all our target GST enzymes (**Fig. S12**), suggesting that GTX binding triggers a global re-structuring of the GST dimer. We performed all comparative HDX-MS measurements shown in **Fig. 7** and **Fig. S8-S11** using this catalytically inert [E:GTX] complex as the reference state to ensure equilibrium binding conditions during the HDX reaction. Below, we highlight overall HDX-MS trends as they relate to catalytic efficiency and the inhibition patterns determined from kinetic studies.

The binding of CDNB to the [E:GTX] complex induced no significant HDX changes in the delta enzymes. Interestingly, CDNB binding to [BmGSTe3:GTX] triggered rigidification of the conserved Asp67-Ser68-His69 motif site in the G-site, as well as in nearly the entire α6 helix, which frames the bottom of the H-site. If a similar structural rigidification occurs in the native [BmGSTe3:GSH:CDNB] complex, it could explain the high catalytic efficiency of BmGSTe3 (**Table 1**). In contrast, CDNB binding to [PxGSTe4:GTX] resulted in a conspicuous loosening in the H-site (helices α4 and α8), as well as in elements of the *N*-terminal domain. These data highlight the structural plasticity of GST enzymes and demonstrate that electrophilic substrate binding (CDNB) may differ between and among GST classes.

ECA binding to the [E:GTX] complexes triggers a rigidification in both Bm enzymes in the G-site D/E-S-H/R motif and α6, and a more variable effect in the Px enzymes that predominantly involves an increase in dynamics. These changes appear to correspond with the trends measured by DSF (**Fig. 6**). ECA binding to the [E:GTX:CDNB] complex triggers organization in all enzymes, but to different extents. Both delta isozymes (which are strongly inhibited by ECA, **Table 2**), undergo a selective rigidification in a contiguous region spanning the C-terminus of α3, the N-terminus of α4, and the intervening loop (Tyr73-Ala93, BmGSTd2 numbering). In addition to the α3/α4 region, ECA binding to BmGSTe3 triggers a much stronger overall rigidification involving the Asp67-Ser68-His69 motif, portions of α1 (including the catalytic Ser11 residue that activates the GSH thiolate) and α8 that, together, frame the H-site, and the conserved Phe100 residue that contributes to the “wafer” motif at the dimer interface^28^. In contrast, the organization of the α3/α4 region in PxGSTe4 is more subdued and is restricted to the *C*-terminal end of α3 and the first few residues of the linker. Curiously, PxGSTe4 was the only enzyme in our set to exhibit non-competitive inhibition with ECA (all other enzymes exhibited a mixed-type pattern). Moreover, all three enzymes that exhibited mixed type inhibition with ECA (BmGSTd2, BmGSTe3, and PxGSTd3) undergo some rigidification of the α3/α4 region, whereas PxGSTe4 undergoes a distinct and characteristic loosening in this structural element.

PER inhibition was slightly more pronounced in the BmGSTs (**Table 2**). Interestingly, there are several distinct changes in the dynamics of the Bm enzymes that are not observed in the Px enzymes upon binding of PER to the [E:GTX:CDNB] complex. First, in the *N*-terminal domain, both Bm enzymes undergo pronounced changes to the α3/α4-linker region, the D/E-S-H/R G-site motif, and the α2/β3-linker G-site motif, which includes the absolutely conserved His residue involved in GSH binding. Additional unique BmGST rigidification is observed in the α4/α5-linker region, which could be involved in α4 helix movements to accommodate ligand binding as observed in the DmGSTe14 crystal structure (**Fig. S5**). In contrast, PxGSTd3 becomes more dynamic in D/E-S-H/R motif, the α6 helix wall of the H-site, and the α1 and β2 elements of the *N-*terminal domain. Finally, PxGSTe4 is rendered less dynamic in the α4 and α8 helices. Therefore, the Bm and Px isozymes undergo distinct conformational changes, upon PER binding.

## DISCUSSION

Our comparative analysis of insect GSTs revealed species-, class-, and isozyme-specific differences in catalytic activity, inhibition, thermal stability, and conformational dynamics that may be exploited for selective insect pest control. Moreover, we found significant variation in the impact of the inhibitors, ECA and PER, on the GSTs. Notably, ECA exhibited substantially stronger inhibition of delta-class GSTs than epsilon-class enzymes. Our data, including HDX and DSF analyses, suggest distinct ECA binding modes or differences in allosteric communication networks between the two classes. In contrast, PER induced predominantly non-competitive inhibition, with substantial species-and isozyme-specific differences.

While ECA binding to delta isozymes resulted in strong mixed-type inhibition, the epsilon-class experienced either mixed-type or non-competitive inhibition (**Fig. 4**). ECA binding induced large increases in the thermal stability of delta enzymes (**Fig. 6**) and very similar, localized rigidification patterns in both delta [E:GTX:CDNB] complexes that differed from the patterns observed in the epsilon enzymes (**Fig. 7**). PxGSTe4 represented a notable outlier across several analyses, displaying non-competitive inhibition and distinct HDX perturbations upon ECA binding. Therefore, this enzyme, or similarly unique GSTs, could be pursued as targets for tailored insecticides against the agricultural pest, *P. xylostella*.

Species and isozyme-specific trends were notably observed for PER inhibition of GST (**Fig. 5**). BmGSTs were inhibited more strongly than Px, and epsilon-class enzymes were more susceptible than the delta-class. Further, BmGSTs were characterized by more extensive organization at the dimer interface and in the α4/α5 linker region upon PER binding. These findings demonstrate that GST inhibitors can exhibit both species and isozyme specificity. Indeed, epsilon-class GSTs from *Anopheles* spp. and other arthropods have been implicated in neutralizing PER, via a speculative mechanism involving binding and sequestration^30^. In *A. aegypti*, partial silencing of epsilon *GST*s resulted in an increased susceptibility to pyrethroid insecticides^8^; however, binding of PER by epsilon-class GSTs was not followed by metabolism or conjugation with GSH. These data showcase the markedly different responses of delta and epsilon GSTs to PER. Widespread resistance to pyrethroids, such as PER, partly via GST-driven metabolism or sequestration, remains a significant concern for disease control.

Significant differences in catalytic efficiency were also observed among GST classes. Epsilon isozymes consistently displayed higher catalytic efficiencies toward the CDNB substrate than delta or sigma enzymes, with BmGSTe3 exhibiting the highest catalytic efficiency among those examined (**Table 1**). Previous studies have shown that elevated CDNB activity can correlate with xenobiotic metabolism and insecticide resistance^29^. Therefore, epsilon GSTs can be regarded as crucial targets for insect pest management. Indeed, the overexpression of *AaGSTe2* and not *AaGSTd1* was responsible for insecticide resistance and elevated metabolism in a DDT and PER-resistant strain of *A. aegypti*^9^.

The exact role of active-site architecture in GST catalysis and its inhibition remains only partially resolved. Epsilon isozymes possess α4-helices inclined approximately 10° further toward the glutathione-binding site than those of delta GSTs (**Fig. 2E,F**, **Table S2**), a key distinguishing feature between the two classes^52^. Closer interactions between the α4-helix and glutathione may promote substrate pre-organization and stabilization of catalytic residues, which could enhance turnover^53^. Furthermore, the displaced *C*-terminal half of this helix alters the H-site creating a more hydrophobic pocket^52^. This displacement may additionally involve restructuring of the α4/α5 linker region (**Fig. S5**). Interestingly, this linker region was rigidified in both *B. mori* [E:GTX:CDNB] complexes with PER, suggesting that α4 helix motion may be an important feature to consider in certain GST inhibition mechanisms. In our HDX-MS analysis, the stabilization of the α4-helix was prominently achieved by GTX binding and further deepened in the Bm isozymes by CDNB and PER co-binding and by CDNB alone in PxGSTe4.

Combinatorial changes in the α4-helix inclination and the α8-helix length may also influence H-site structure, substrate recognition, and inhibitor access^38,52^. For example, the α8-helix in *Anopheles gambiae* AgGSTe2 contains 10 more residues than its delta counterparts, pulling α8 further from the active site and purportedly creating a larger substrate entry gate^38^. Furthermore, the *C*-terminal extension can interact with residues in the α4/α5 linker noted above^52^, which, in turn, could also affect the entrance of ECA/PER molecules. Our HDX-MS analyses revealed highly variable ligand-dependent alterations to α8-helix dynamics. For example, in BmGSTd2, the entire α8 became rigid upon PER binding, while the opposite was observed for PxGSTd3 (**Fig. 7**). The speculated impact of the α8-helix on inhibitor access would have to be further investigated, with a larger cohort of enzymes, to draw strong conclusions. Finally, the α6 helix, which forms one wall of the H-site, was observed to rigidify upon ligand binding in our HDX studies. This structuring likely involves a strongly conserved Ser residue in α6, which H-bonds to the highly conserved His/Arg residue of the D/E-S-H/R G-site motif^27^. Thus, the α6 element and its organization may be structurally linked to dynamics at the core of the dimer in some GST enzymes.

A broader finding of this study is the remarkable diversity in conformational dynamics among insect GSTs and the absence of universal structural signatures associated with inhibition. Although some relationships between ligand-induced dynamics and inhibitory mechanisms were apparent, GSTs likely require substantial conformational flexibility to accommodate the wide range of electrophilic substrates encountered during detoxification. Consequently, inhibitor binding modes, allosteric networks, and regulatory mechanisms are expected to vary considerably across this enzyme family. Continued systematic biochemical and biophysical investigations will therefore be necessary to better understand GST structure-function relationships and to exploit these differences for the development of selective insecticides.

## MATERIALS AND METHODS

### Plasmids and *E. coli* strains

The plasmid vector used for recombinant protein purification in *E. coli* was pET28α (+) (Invitrogen, MA, US). The *E. coli* strain NEB10β was purchased from New England Biolabs (NEB, MA, US) and was used for plasmid accumulation. The *E. coli* strain BL21 (DE3) was used for recombinant protein expression.

### Chemicals

Analytical and assay-grade standards were purchased for enzyme kinetics and inhibition assays, including reduced glutathione (GSH) (98%; Oakwood chemicals SC, US, Cat. 213215), 1-chloro-2,4-dinitrobenzene (CDNB) (97%; Sigma Aldrich, MO, US, Cat. 138630), ethacrynic acid (≥98%; NY, US, Cat. BML EI128), permethrin (VWR, PA, US, Cat. 89161-758) and *S*-hexylglutathione (GTX) (>98%, Millipore Sigma, MA, US, Cat. H6886).

### Structural and network analysis

The amino acid sequence of BmGSTe3 was used to generate a structural model using an AlphaFold pipeline^54^. To assess structural conservation and visualize potential ligand-binding sites, the BmGSTe3 model was structurally aligned with the crystal structure of *Drosophila melanogaster* epsilon GST (PDB ID: 7DB4), which includes co-crystallized ligands; glutathione (GSH) and H2C (IUPAC name: 1-(4-fluorobenzyl)-1H-thieno[3,2-c][1,2]thiazin-4(3H)-one 2,2-dioxide). *Anopheles gambiae* delta and epsilon GSTs (PDB IDs: 1PN9 and 2IL3, respectively) were included as references due to the availability of their crystal structures. The alignment was visualized with ESPript 3.0^55^, where secondary structure elements were annotated based on the reference crystal structures. The inclination angle of the α4 helix toward the GSH-binding site was calculated in PyMOL by generating centroid pseudoatoms from the C atoms of the lower (residues 108-111) and upper (residues 118-121) portions of the α4 helix after alignment to AgGSTe2 (PDB: 2IMK), and computing the angle between the helix axis vector and the vector from the helix tip toward the ligand centroid. A smaller angle indicates greater tilt of the α4 helix towards the substrate-binding pocket. The construction of the GST sequence similarity network using the Enzyme Function Initiative webtool is described in the main text^35,56^.

### Construct design and assembly

Full-length genes encoding delta (d), epsilon (e), and sigma (s) class glutathione *S*-transferases (GSTs) from *Aedes aegypti* (Aa), *Apis mellifera* (Ap), *Bombyx mori* (Bm), *Leptinotarsa decemlineata* (Ld), and *Plutella xylostella* (Px) were retrieved from the NCBI database and synthesized by Twist Biosciences (CA, US). The following gene sequences were codon-optimized for expression in *E. coli*: *ApGSTd1* (NM_001358979.1), *BmGSTd2* (AB176691), *LdGSTd3* (KU522308), *PxGSTd3* (AB541016), *AaGSTe2* (AY819710), *BmGSTe3* (EF506488), *PxGSTe4* (KF929205), and *BmGSTs1* (AB206971). Each gene was synthesized with *BamHI* and *XhoI* restriction sites at the 5′ and 3′ ends, respectively. Plasmid DNA (1 μg of pET28α(+) vector and 750 ng of insert) was digested with *BamHI* and *XhoI* (New England Biolabs (NEB), MA, US) at 37 °C for 1 hour. Digestion was confirmed by agarose gel electrophoresis (1% w/v) using 3 μL of Gel Loading Dye, Purple (6×; NEB). The linearized vector and digested GST inserts were purified using the Monarch® Gel Extraction Kit (NEB). Ligation was performed overnight at 4 °C using a 3:1 molar ratio of insert to pET28α(+) vector with 1 μL of T4 DNA Ligase (NEB). A 5 μL volume of the ligation reaction was transformed into 50 μL of chemically competent *E. coli* NEB10β cells. Transformants were selected on LB agar plates containing 50 μg/mL kanamycin and incubated overnight at 37 °C. Single colonies were screened by colony PCR using 2× Quick-Load Taq polymerase (NEB), with both T7 sequencing primers and gene-specific primers. An empty pET28α(+) plasmid served as a negative control. A complete list of primers used in this study is provided in **Table S3**. Plasmids from PCR-verified colonies were isolated using the EZ-10 Spin Column Plasmid DNA Miniprep Kit (Bio Basic Inc., ON, Canada). For sequence confirmation, positive clones were sequenced for whole plasmid sequencing at Plasmidsaurus (KY, US).

### Recombinant protein production

Recombinant protein expression was initiated by culturing *E. coli* BL21 (DE3) in LB media at 37 °C and 225 rpm until mid-log phase (OD_600_ of 0.5). Protein expression was induced by adding 1OmM isopropyl β-D-1-thiogalactopyranoside (IPTG; VWR International, PA, US), followed by incubation at 25 °C for 4Oh. Cells were harvested by centrifugation at 1,000O×Og and resuspended at a 1:10 (w/v) ratio in buffer A (20OmM Tris-HCl, pHO8.0, 300OmM NaCl, 10% (v/v) glycerol). Cell lysis was performed on ice via sonication using a microprobe ultrasonic processor (Misonix, NY, US) with cycles of 5Os, on, and 5Os, off, for a total of 5Omin at 80OHz. The lysate was clarified by centrifugation at 14,000O×Og for 30Omin to separate the soluble fraction from cell debris. His-tagged proteins were purified using TALON® Superflow™ metal affinity resin (Cytiva, Sweden) via gravity-flow method. The clarified lysate was incubated with the resin for 1Oh at 4 °C under gentle agitation, then transferred to a disposable polypropylene column (Thermo Fisher Scientific, MA, US). The column was washed with 5 bed volumes of buffer A supplemented with 5OmM imidazole, and target proteins were eluted with 10 bed volumes of buffer B (20OmM Tris-HCl, pHO8.0, 300OmM NaCl, 150OmM imidazole, 10% (v/v) glycerol). The elute was concentrated using Vivaspin® centrifugal filter units (10OkDa MWCO; Sartorius, Germany). Protein concentrations were determined by the Bradford assay (Bio-Rad, CA, US) using bovine serum albumin as the standard. The purified GST proteins were assessed by SDS-PAGE as well. Pure proteins were then diluted to 1 mg/mL aliquots and flash-frozen in liquid nitrogen and stored at-80 °C until further use.

### *In vitro* kinetic assays

Enzymatic activity with the general GST substrate, 1-chloro-2,4-dinitrobenzene (CDNB), was measured with minor modifications to established protocols^39^. Steady state kinetic assays were conducted for selected GSTs by varying CDNB concentrations from 50-3200 μM, while maintaining a constant concentration of GSH at 5 mM. Reactions were performed in triplicate using 200 ng of enzyme per well in 1× PBS buffer (pH 6.5), using flat-bottom, high-binding 96-well microplates (Sarstedt, Germany). Kinetic measurements were acquired using a SpectraMax M3 plate reader (Molecular Devices, CA, US) in kinetic mode, by monitoring the increase in absorbance at 340 nm due to the formation of the CDNB conjugation to the thiol groups of glutathione (ε_340_ = 9.6 mM⁻¹·cm⁻¹). Negative controls without enzyme were included, and all assays were performed in triplicate. The initial velocities of the reactions were calculated from the absorption change (ΔA340) prior to the consumption of the first 10% for the CDNB starting material. Inhibition assays were conducted using the well-established GST inhibitors permethrin (PER), a pyrethroid insecticide, and ethacrynic acid (ECA). Reactions were performed in triplicate at a saturating GSH concentration (5 mM) with CDNB as the variable substrate, in the presence of 0, 1, 10, 25, and 50 μM of either PER or ECA. Both compounds were dissolved in ethanol; therefore, all assay conditions, including controls, contained a consistent ethanol concentration to eliminate solvent-related variability. All kinetic data were fitted with non-linear regression to the Michaelis-Menten equation using OriginPro (OriginLab, MA, US). The maximum inhibition (as reported in **Table 2**) was estimated for each enzyme/inhibitor pair by fitting the inhibitor concentration dependence of the steady state kinetic parameters to hyperbolic or linear functions.

### Thermal shift assays

Effect of ligand binding on melting temperature (Tm) of GST was measured with 0, 0.049, 0.390, 3.130, 50, and 100 μM of PER and ECA in the presence of fixed concentration of GSH (2 mM). Each enzyme was also tested with GTX, a hydrophobic analog of GSH, either alone or in combination with CDNB, PER, or ECA, 5 μM of each protein, 10X SYPRO Orange dye (Sigma Aldrich) and 50 mM HEPES (pH 7.5) were mixed with ligand. Protein, buffer, and ligand were incubated on ice for 20 minutes, then 10X SYPRO dye was added at the end. A control with no protein was also added. Protein melting temperatures were assayed using the Bio-Rad family of CFX Real-Time PCR Systems using the following conditions: a continuous increase from 10.0 °C to 95 °C was scanned every 0.5 °C for 10 seconds. Change in melting temperature (ΔTm) was determined by subtracting the melting temperature of the unbound protein from that of the ligand-bound protein. Mean ΔTm was plotted against ligand concentration for each protein using data from three replicates.

### Hydrogen-deuterium exchange (HDX) reactions

HDX-MS experiments were performed to investigate ligand-induced conformational changes in four selected GSTs: BmGSTe3, BmGSTd2, PxGSTe4, and PxGSTd3. All exchange reactions were carried out in 1X PBS buffer (pH 6.5) prepared in H₂O. Final reaction mixtures (50 µL) contained 2 μM of the respective GST protein and, where applicable, 2 mM GSH, 2 mM GTX, 3.2 mM CDNB, 50 μM PER, and ECA. The eight ligand-binding states analyzed in detail were: GST alone; [E:GSH], [E:GTX]; [E:GTX:CDNB]; [E:GTX:PER]; [E:GTX:ECA], [E:GTX:CDNB:PER] and [E:GTX:CDNB:ECA]. Ligands were prepared in their respective solvents: GTX and ECA in DMSO, CDNB and PER in ethanol. To maintain consistent solvent conditions across all samples, equal volumes of ethanol and DMSO were added to each reaction, keeping the total organic solvent concentration at approximately 2% (v/v). Exchange reactions were initiated by diluting enzyme to a final concentration of 2 μM in a 50 μL reaction volume containing 70% D_2_O. After 5 minutes of deuterium labeling at room temperature, reactions were quenched by addition of 50 μL ice-cold quench buffer (1 M guanidine-HCl, 100 mM phosphate, pH 2.0). Quenched samples were immediately flash frozen in liquid nitrogen and stored at-80 °C until mass spectrometry analysis. All HDX reactions were performed on the same day in triplicate using the same stock solutions to ensure reproducibility. HDX-MS experiments were performed on a Waters Synapt G2-Si mass spectrometer equipped with HDX technology and maintained at 0.4 °C. Samples were thawed at 35 °C for 60 s and injected into a 40 μL loop exactly 2 min after removal from-80 °C. Data analysis was performed using ProteinLynx Global SERVER (PLGS, Waters, MA, US) and DynamX 3.0 (Waters, MA, US) as described previously^57^. Statistical significance of HDX uptake differences were determined with Deuteros 2.0^58^.

## STATISTICS AND REPRODUCIBILITY

All experiments were conducted on a minimum of *n*□=□3 biologically independent samples and analyzed by a two-tailed, unpaired Student’s *t*-test. To compare the significance of all groups within a set of assays, the data were analyzed using one-way analysis of variance (ANOVA) with subsequent pairwise comparisons conducted using Tukey’s HSD test (GraphPad v.10.5.0). Data is represented as sample mean ± standard error of the mean. Statistical significance was determined as *p <* 0.05 and is denoted by letters. Letters denote statistically similar groups between and within treatments.

## DATA AVAILABILITY

The authors declare that the data supporting the findings of this study are available within the paper and its **Supplementary Information** files. Source data are provided within this paper.

## ACKNOWLEDGEMENTS

This work was funded by the New Frontiers in Research Fund – Exploration (NFRF-E) (grant number: NFRFE-2022-00316), entitled “Plant-derived biosynergists to enhance pesticide efficacy.” Dr. Sharma reports a postdoctoral award from the Centre for Structural Biology (CRBS) at McGill University.

## COMPETING INTERESTS

The authors declare no competing interests.

## AUTHOR CONTRIBUTIONS

MS prepared the first draft of the manuscript. MS devised and conducted experiments, including structural analysis, protein expression and purification, enzyme kinetics, enzyme-inhibition assays, and DSF. MS and SQ jointly conducted HDX-MS experiments, with conceptual and technical guidance by CJT. All co-authors contributed to analysis and interpretation of data. MD and CJT conceived the study and acquired the funding. CJT supervised SQ and revised the manuscript. MD and CJT guided the study, supervised MS, revised the manuscript, and approved it for submission.

## SUPPLEMENTARY FIGURES

**Fig. S1.**
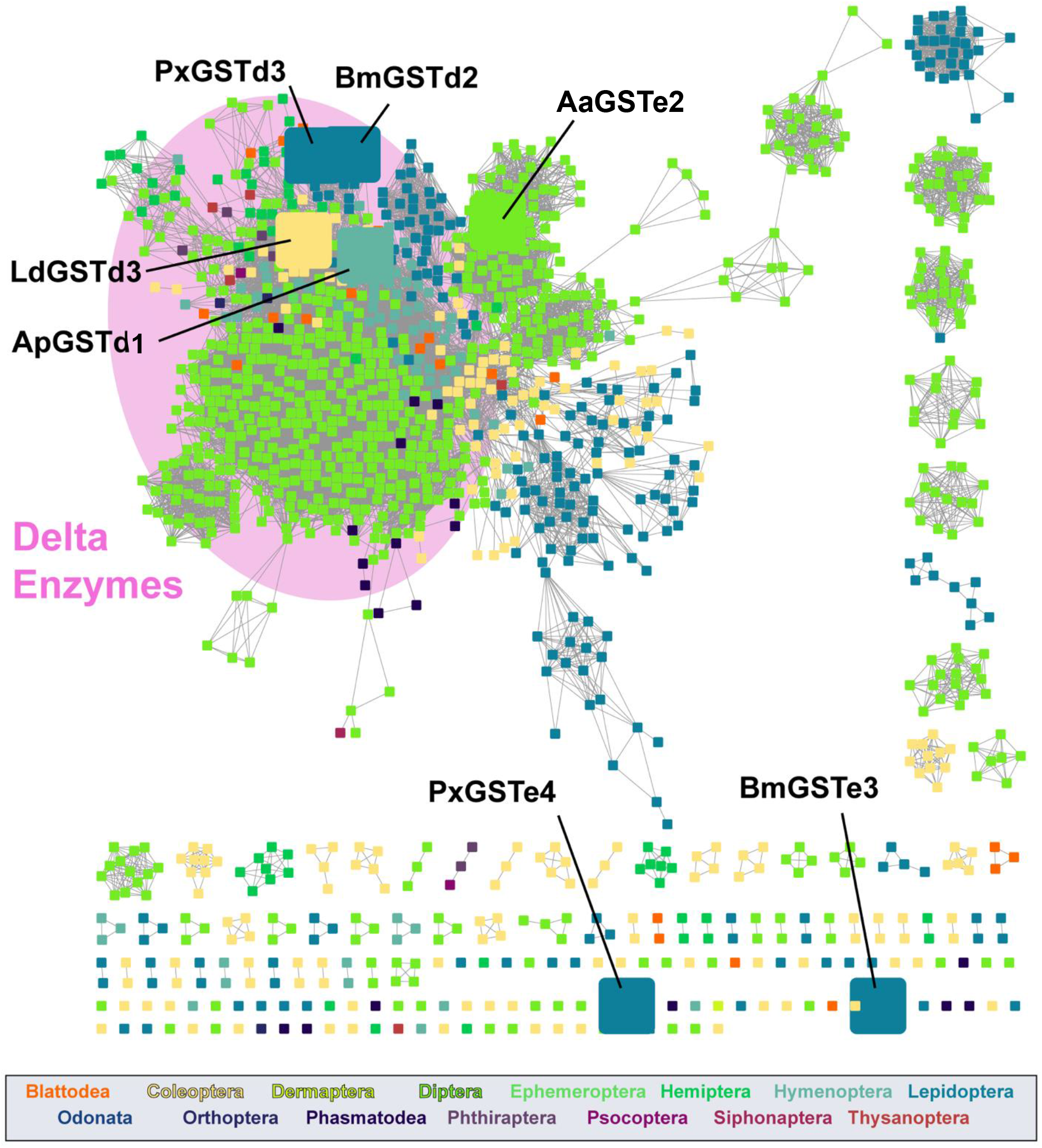
Sequence similarity network of insect delta and epsilon GST enzymes. The delta and epsilon enzymes from the full network (Fig. 1, alignment score *E* = 65) are displayed here at a higher alignment score (*E* = 72) to better illustrate the closer sequence similarity (higher clustering) among the delta enzymes and the higher sequence divergence among the epsilon enzymes. The nodes corresponding to our target enzymes are enlarged.

**Fig. S2.**
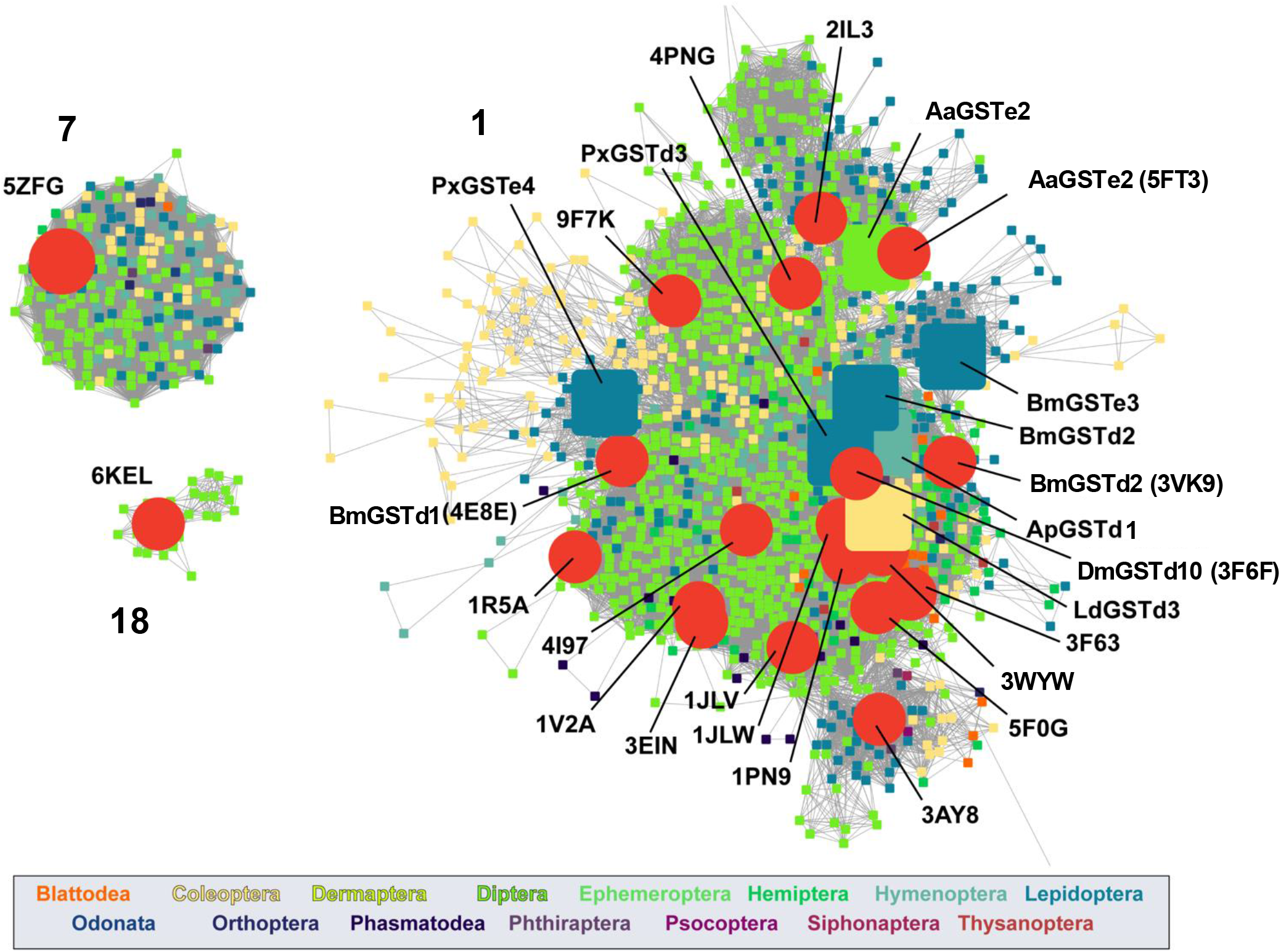
**Sequence similarity relationships between target GST enzymes and enzymes with solved high-resolution structures**. Network clusters are numbered as in Fig. 1a. The PDB IDs for enzymes with solved structures are indicated.

**Fig. S3.**
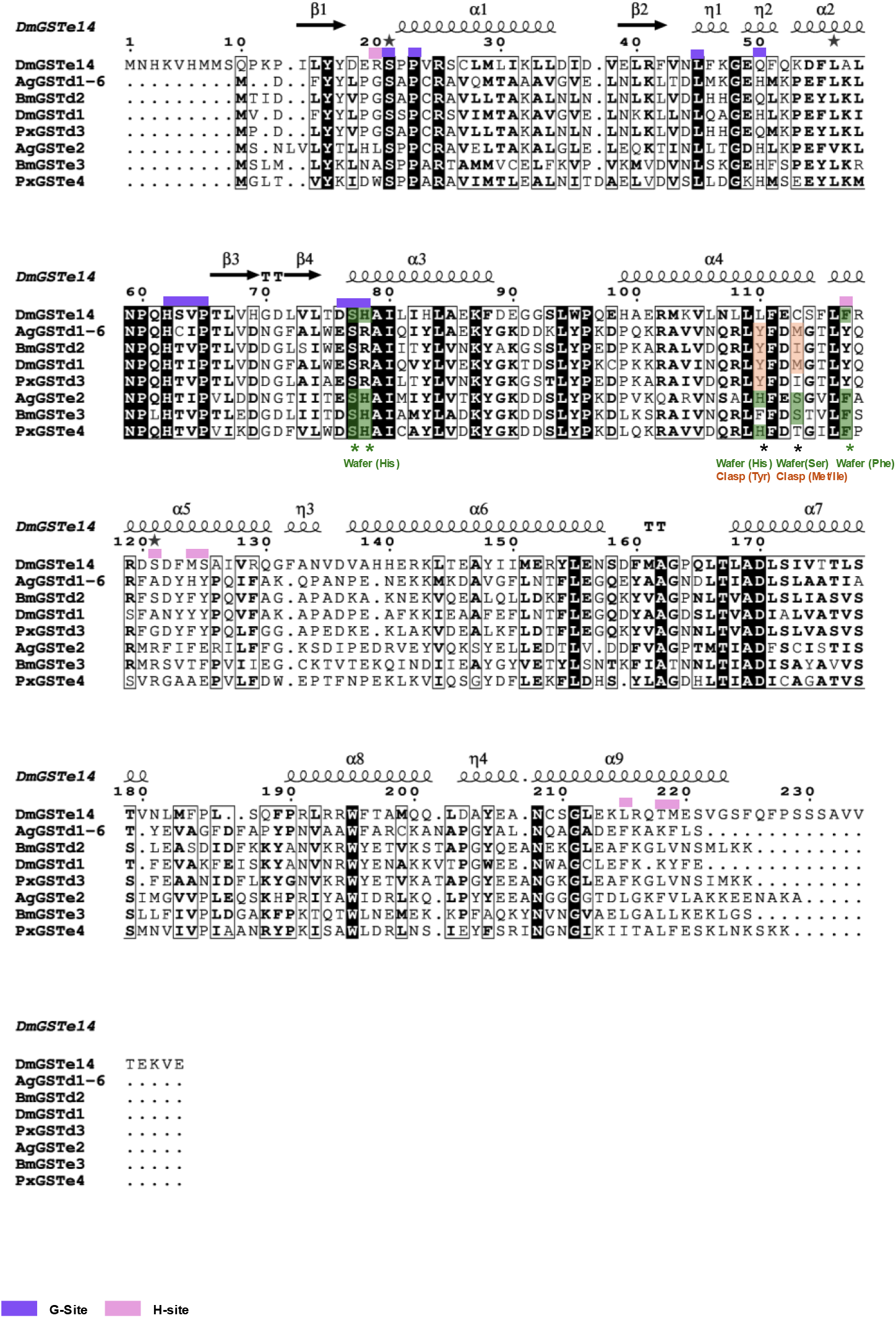
Amino acid sequence alignment of selected GSTs. Identical sequences are marked in black background/white text. The G-site and H-site residues are indicated by purple and pink bars above the alignment, respectively, based on the crystal structures of DmGSTe14 (PDB ID: 7DB4). Asterisks mark residues conserved across all species. Secondary structure elements: α-helices (α) and β-strands (β) are annotated according to the reference crystal structures. Ag: *Anopheles gambiae*; Bm: *Bombyx mori*; Dm: *Drosophila melanogaster*; Ld: *Leptinotarsa decemlineata*; Px: *Plutella xylostella*. d: delta and e: epsilon GST.

**Fig. S4.**
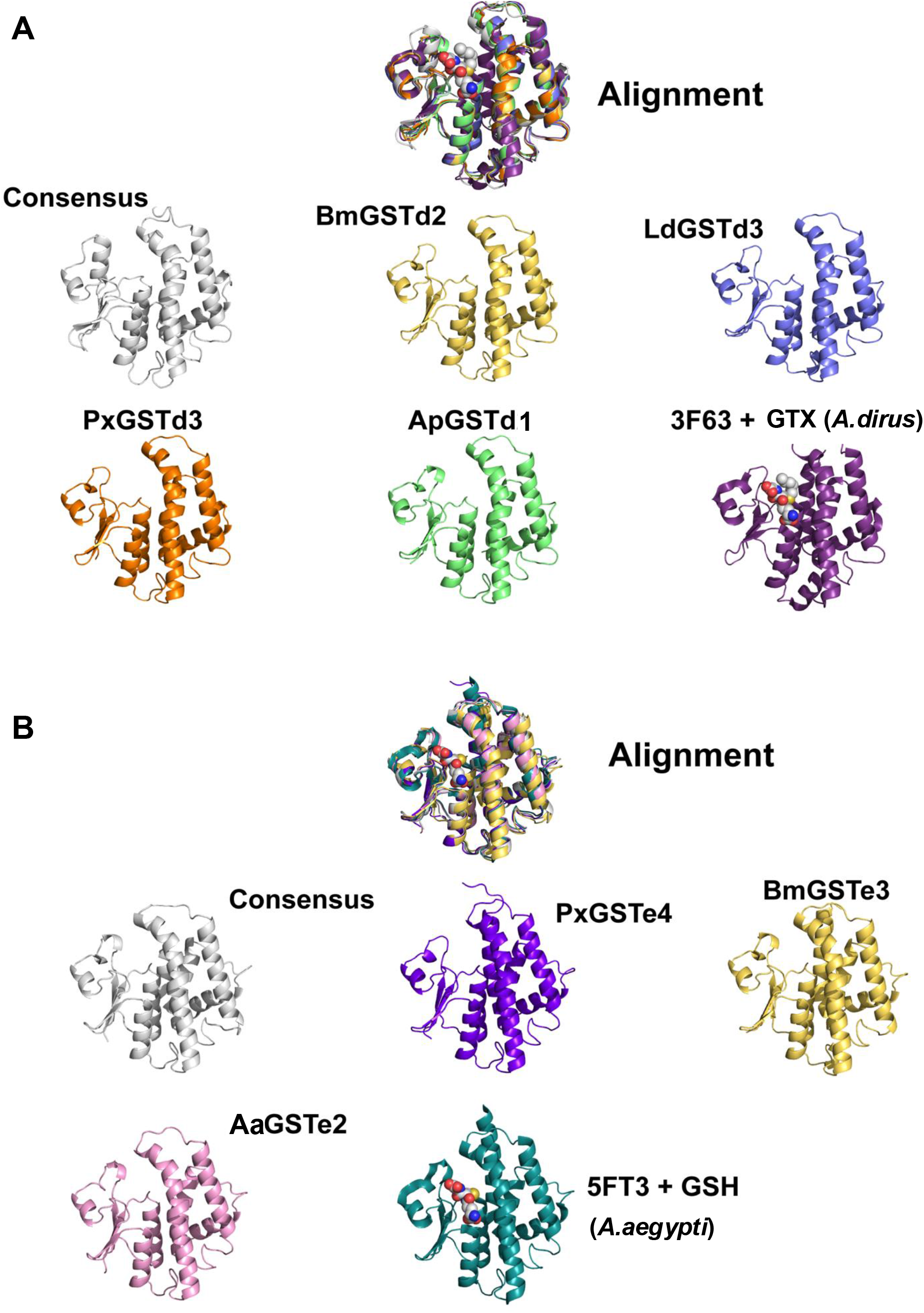
Structural alignments of target GST enzymes. AF3 models of target GST enzymes were aligned with (i) the AF3 model of the consensus structure derived from the sequence alignment of enzymes in the sequence similarity network (Fig. 1A) and (ii) with a ligand-bound experimental structure of a closely related enzyme (see **Fig. S2**). In general, these alignments suggest high accuracy of the AF3 models of our target enzymes.

**Fig. S5.**
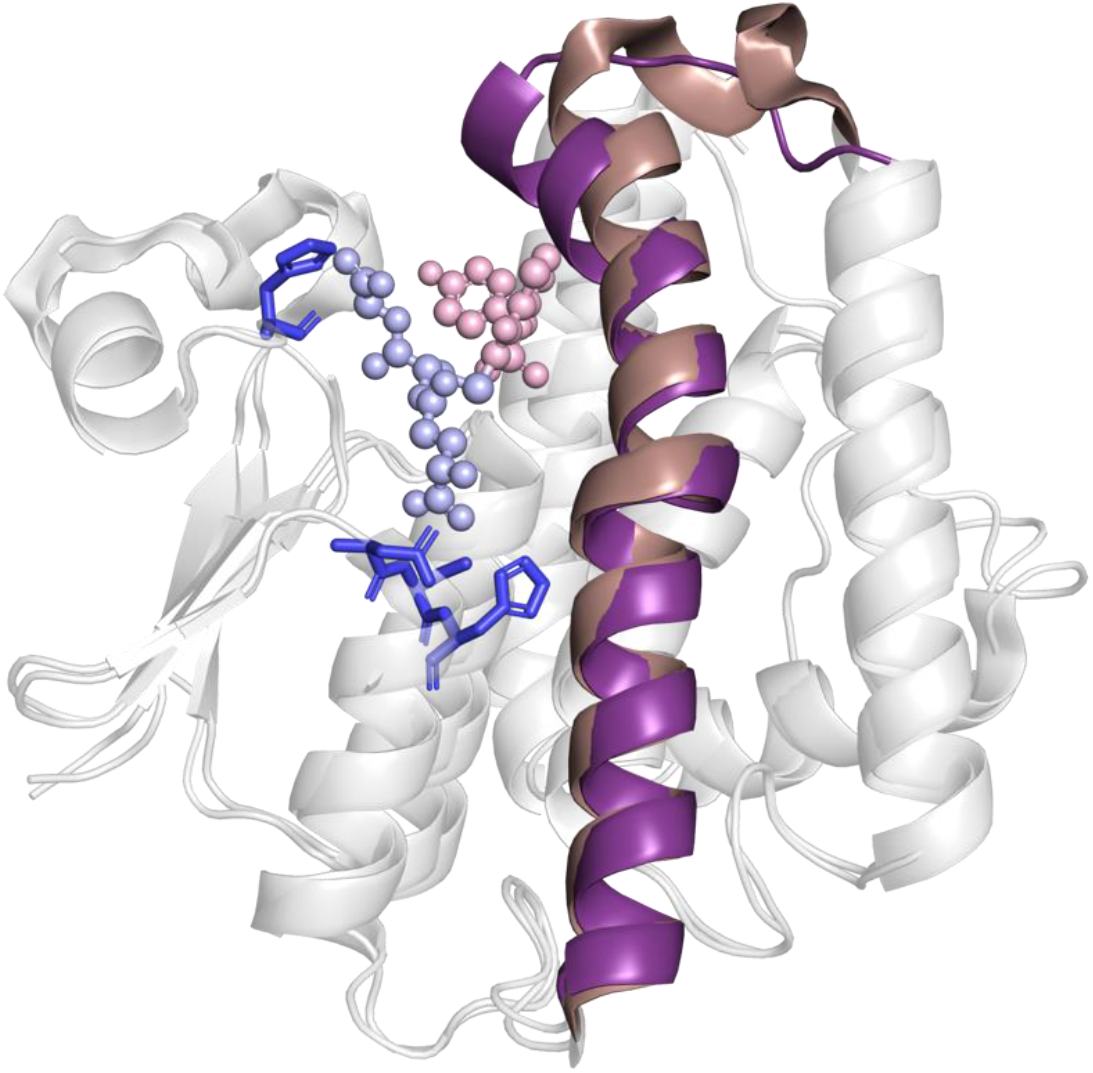
Structural alignment of BmGSTe3 and *Drosophila melanogaster* epsilon GST (PDB ID: 7DB4), featuring their α4-helices, coloured in violet and dark salmon, respectively; GSH ligand is shown in light blue and the ligand H2C shown in light pink against the grey enzyme.

**Fig. S6.**
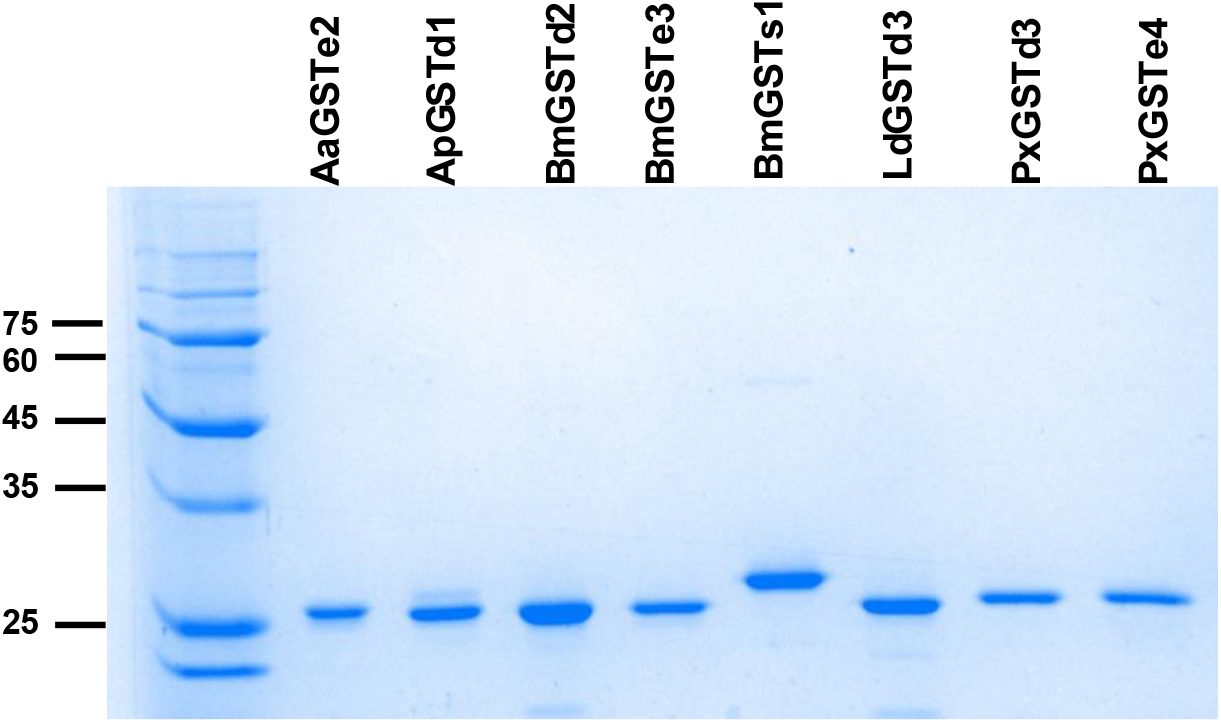
Purified recombinant proteins for *in vitro* assays. Purified proteins (700 ng) were observed with Coomassie Brilliant Blue R250 staining. Expected molecular masses of proteins are: AaGSTe2, 28.23 kDa; ApGSTd1, 28.71 kDa; BmGSTd2, 27.77 kDa; BmGSTe3, 27.90 kDa; BmGSTs1, 26.88 kDa; LdGSTd3, 28.03 kDa; PxGSTd3, 27.46 kDa; PxGSTe4, 28.30 kDa

**Fig. S7.**
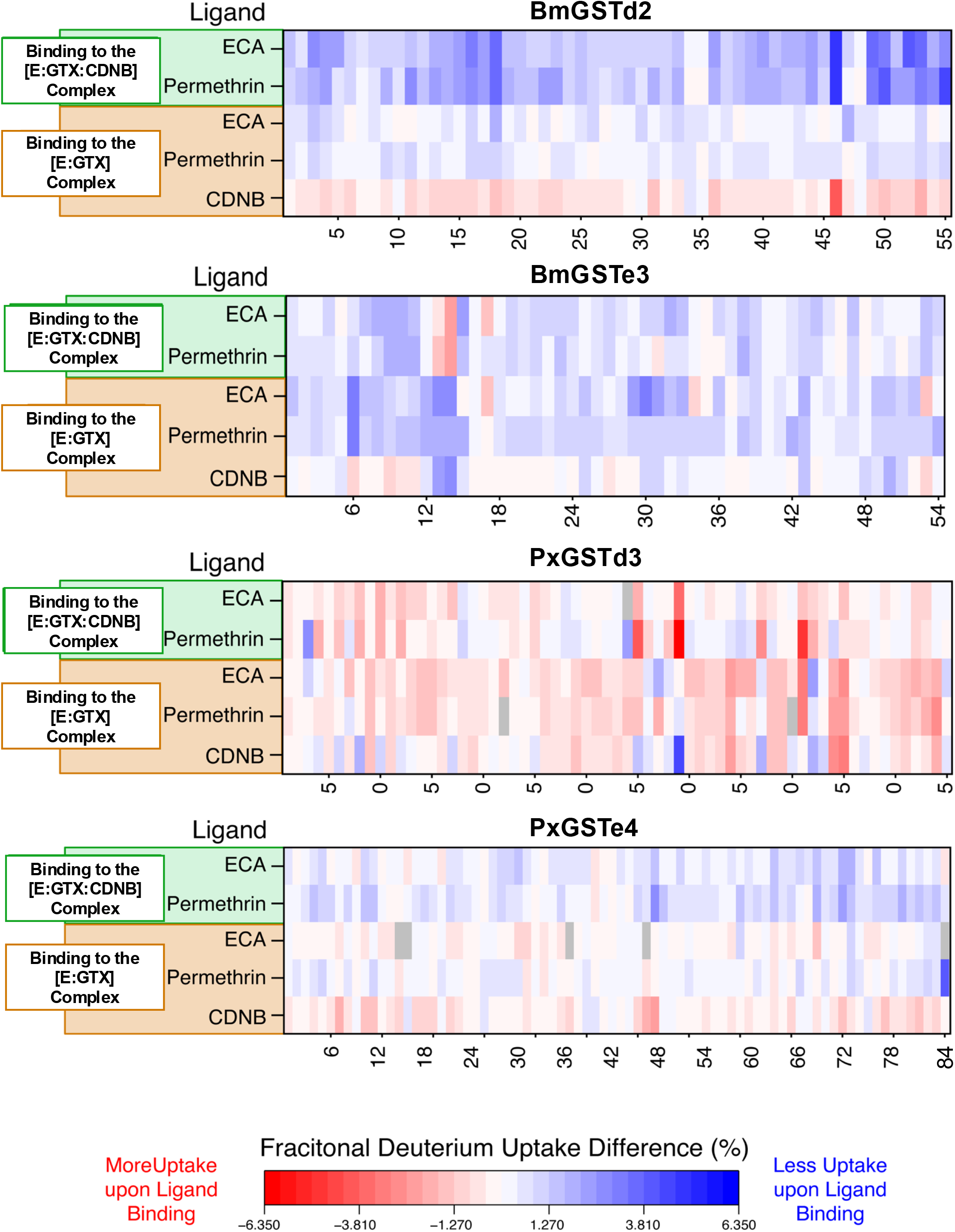
Comparison of deuterium uptake differences. Hydrogen-deuterium exchange (HDX) data from Fig. 7 of the main text are re-plotted in heat map format. The heatmaps show the fractional deuterium uptake differences (% change in peptide deuterium level) between [E:GTX] and other ligand-bound states ([E:GTX:CDNB], [E:GTX:ECA], [E:GTX:ECA:CDNB], [E:GTX:PER], and [E:GTX:PER:CDNB]) for the target enzymes. Blue/red regions represent decreased/increased deuterium uptake, respectively, according to the difference calculations shown to the left of each panel. All maps are coloured on the same scale to allow visualization of relative HDX changes between enzymes. For each enzyme, HDX peptides are arranged from left to right along the x-axis in an *N*-to *C*-terminal direction. A complete, numbered inventory of every HDX peptide for each heat map is provided as supporting data. Regions with a significant uptake difference (*p* < 0.01) are coloured similarly on the AF3 model structures in Fig. 7.

**Fig. S8.**
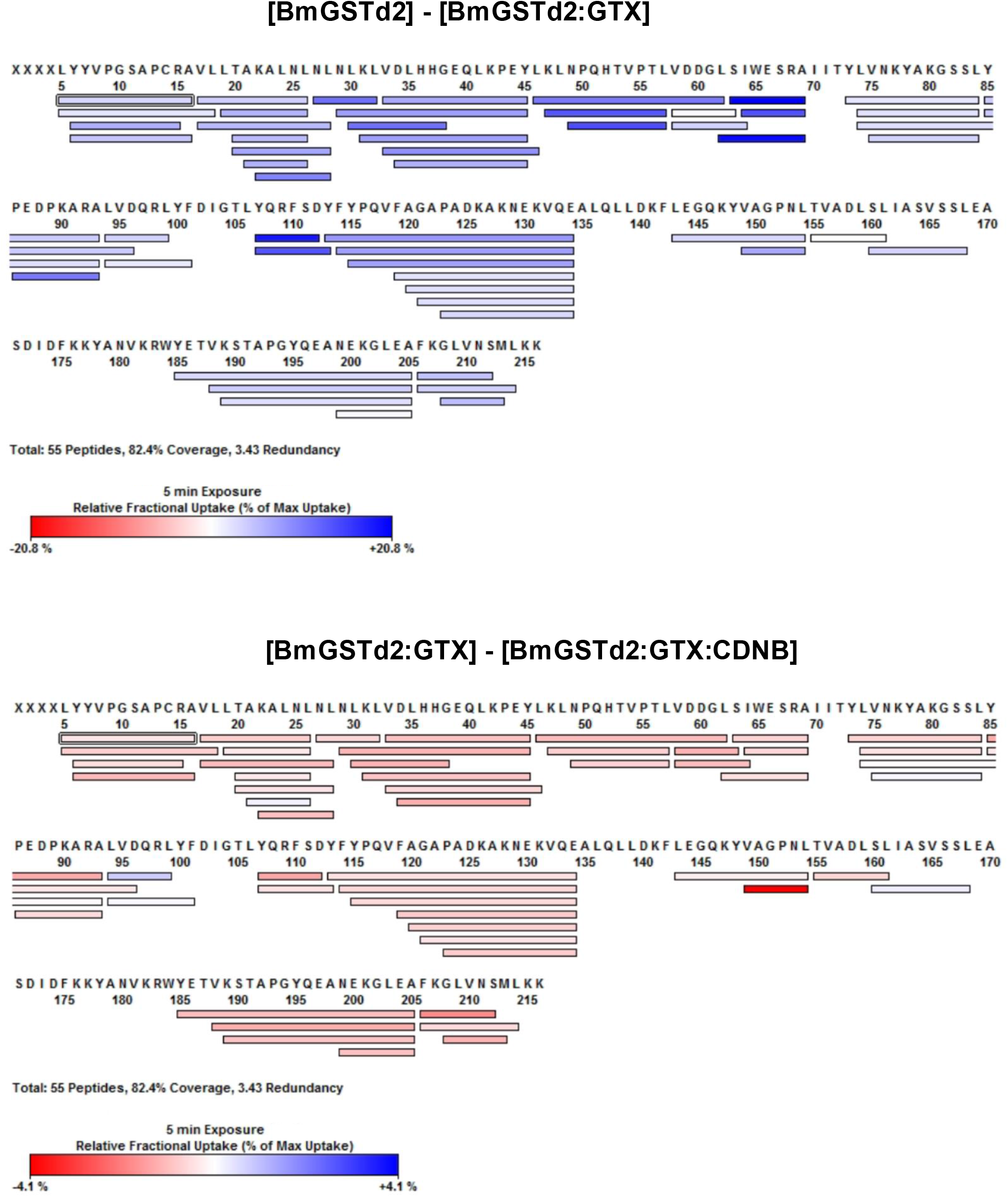
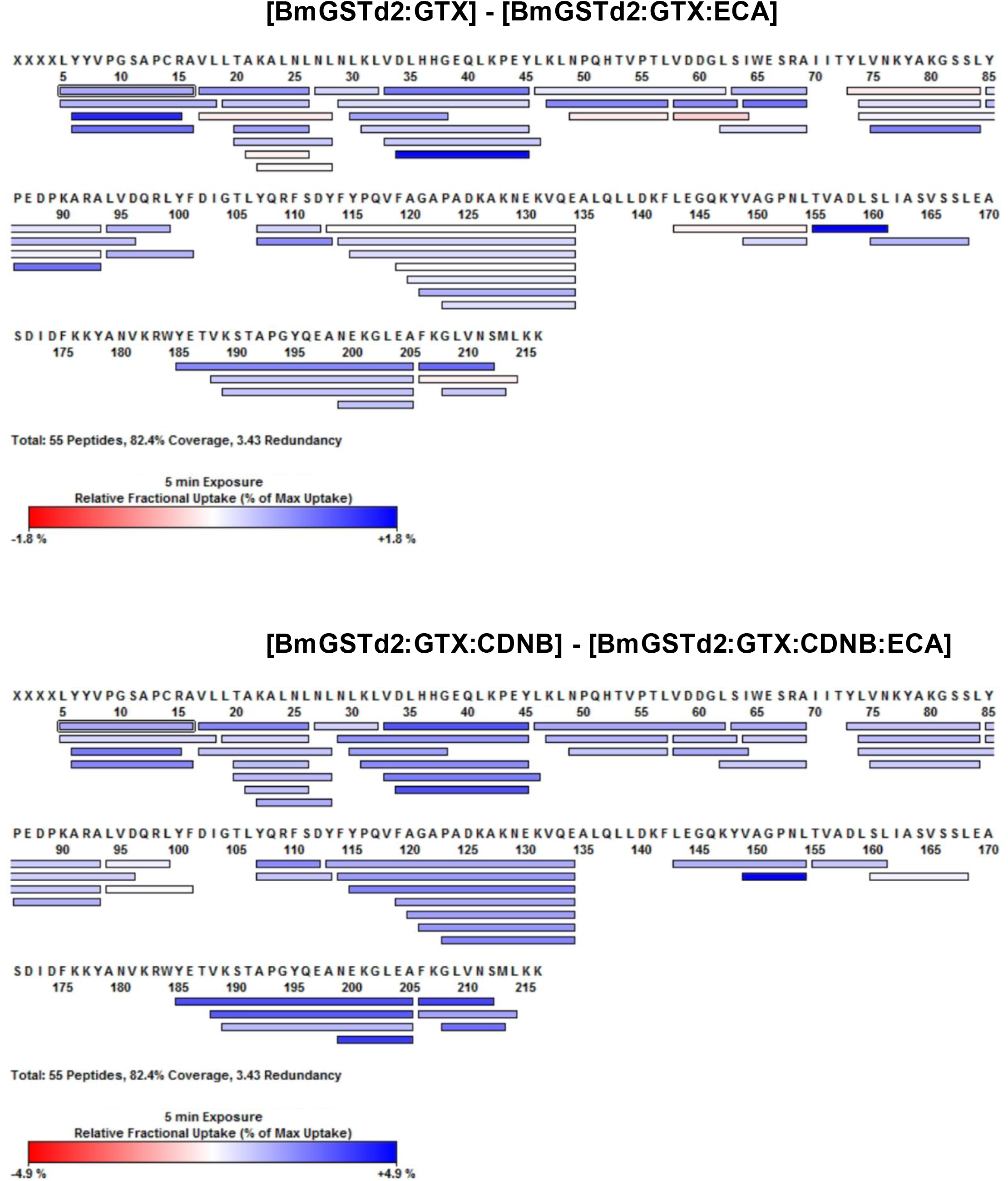
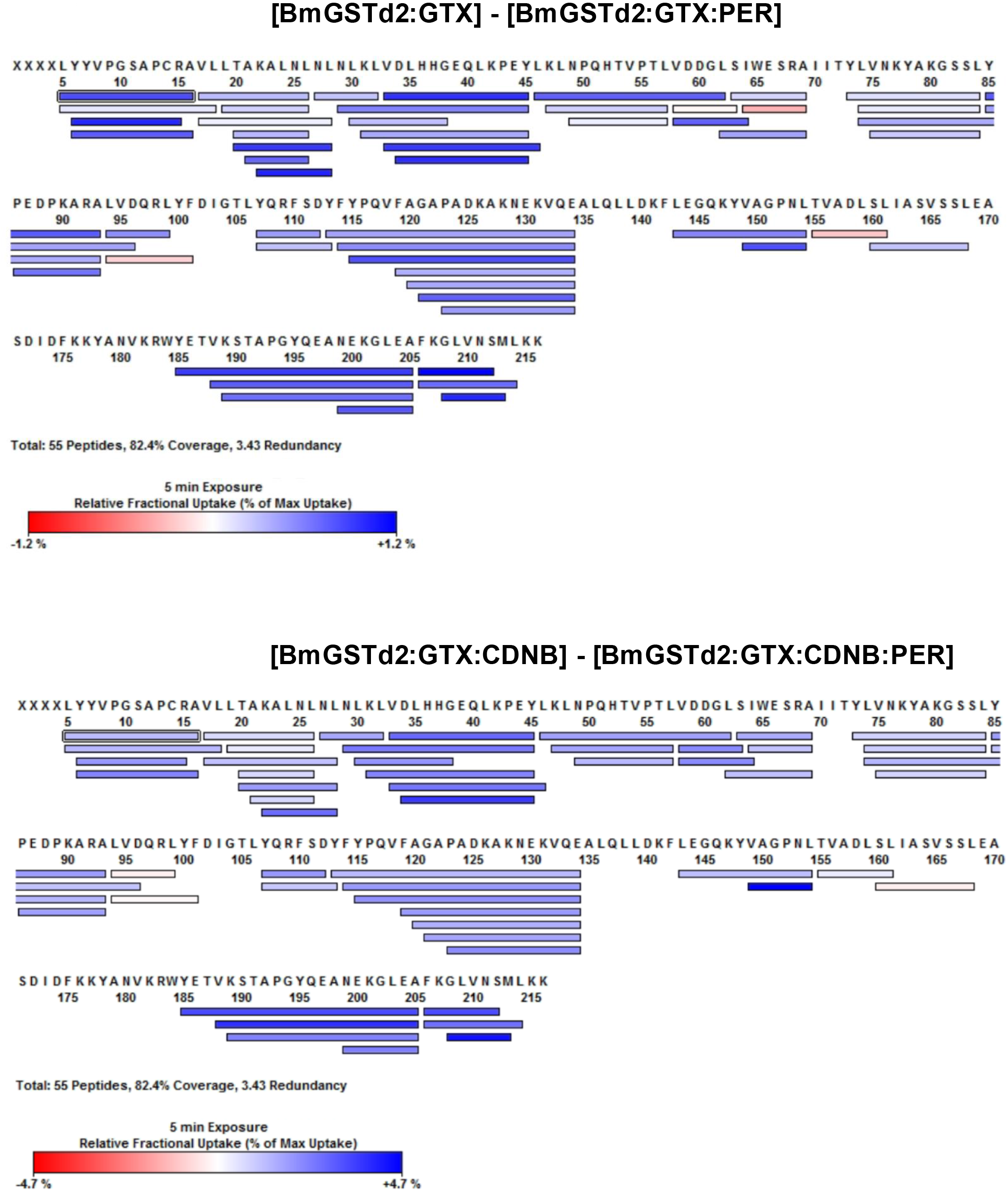
**Coverage maps of HDX-MS for individual states for BmGSTd2**

**Fig. S9.**
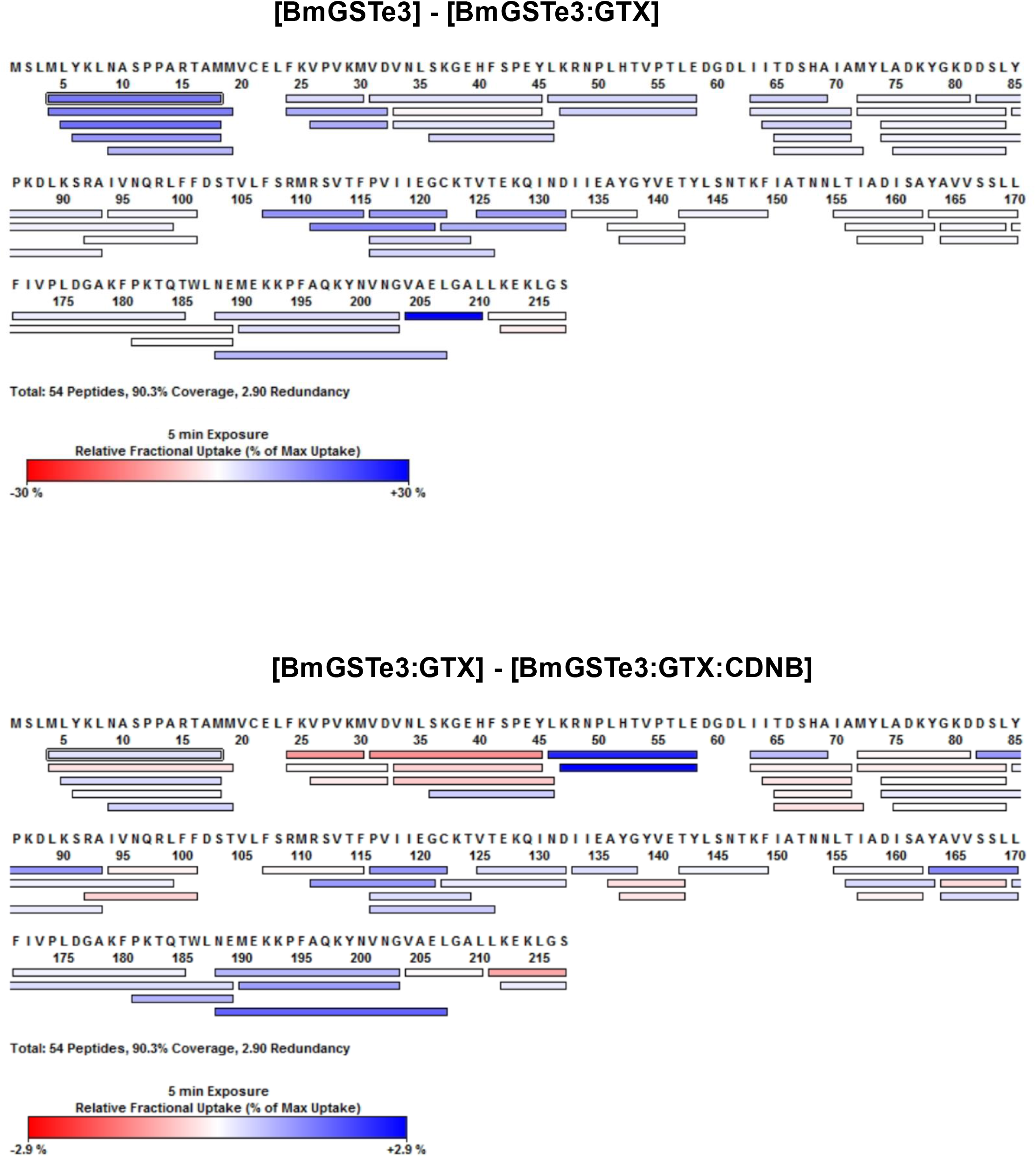
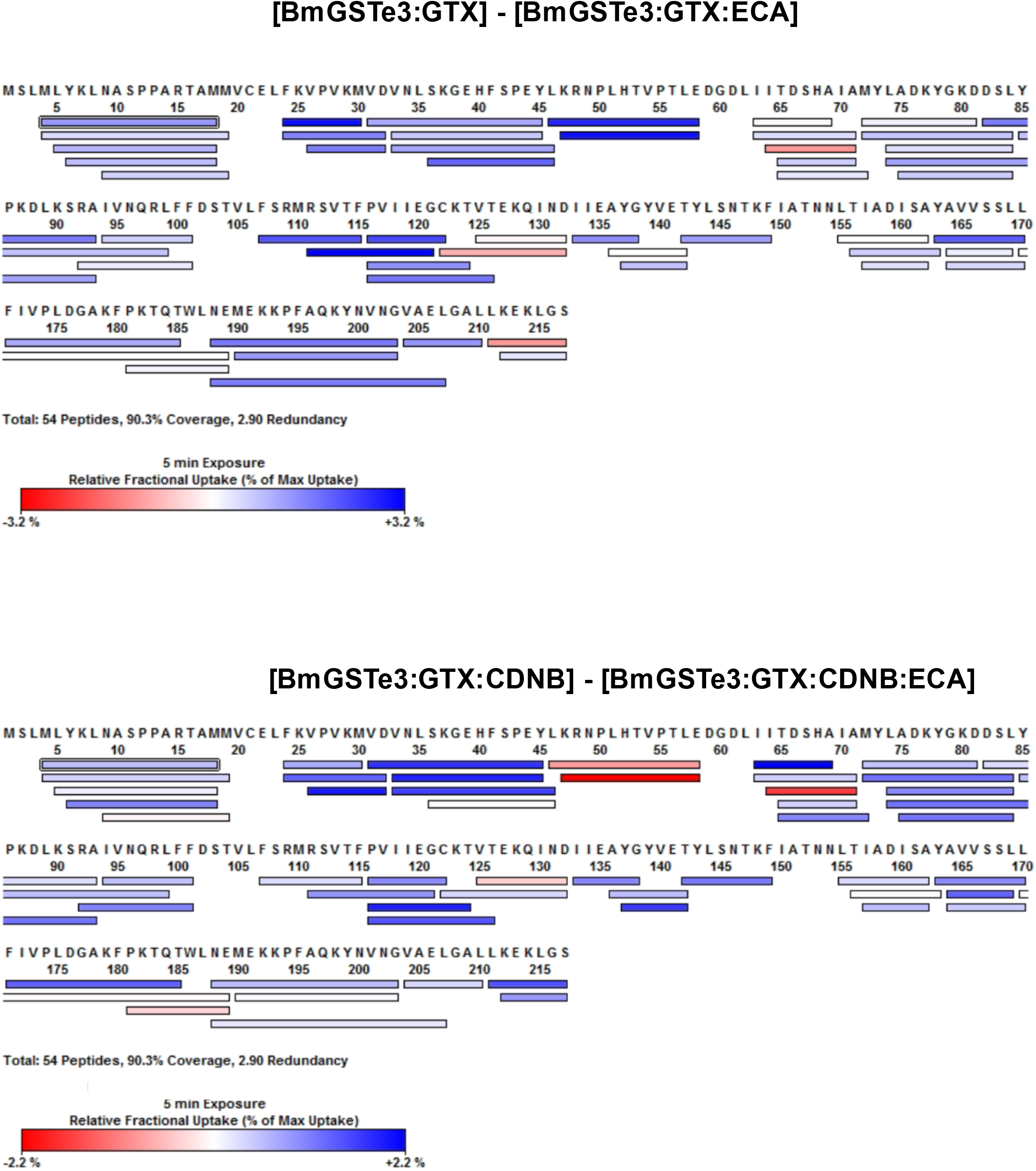
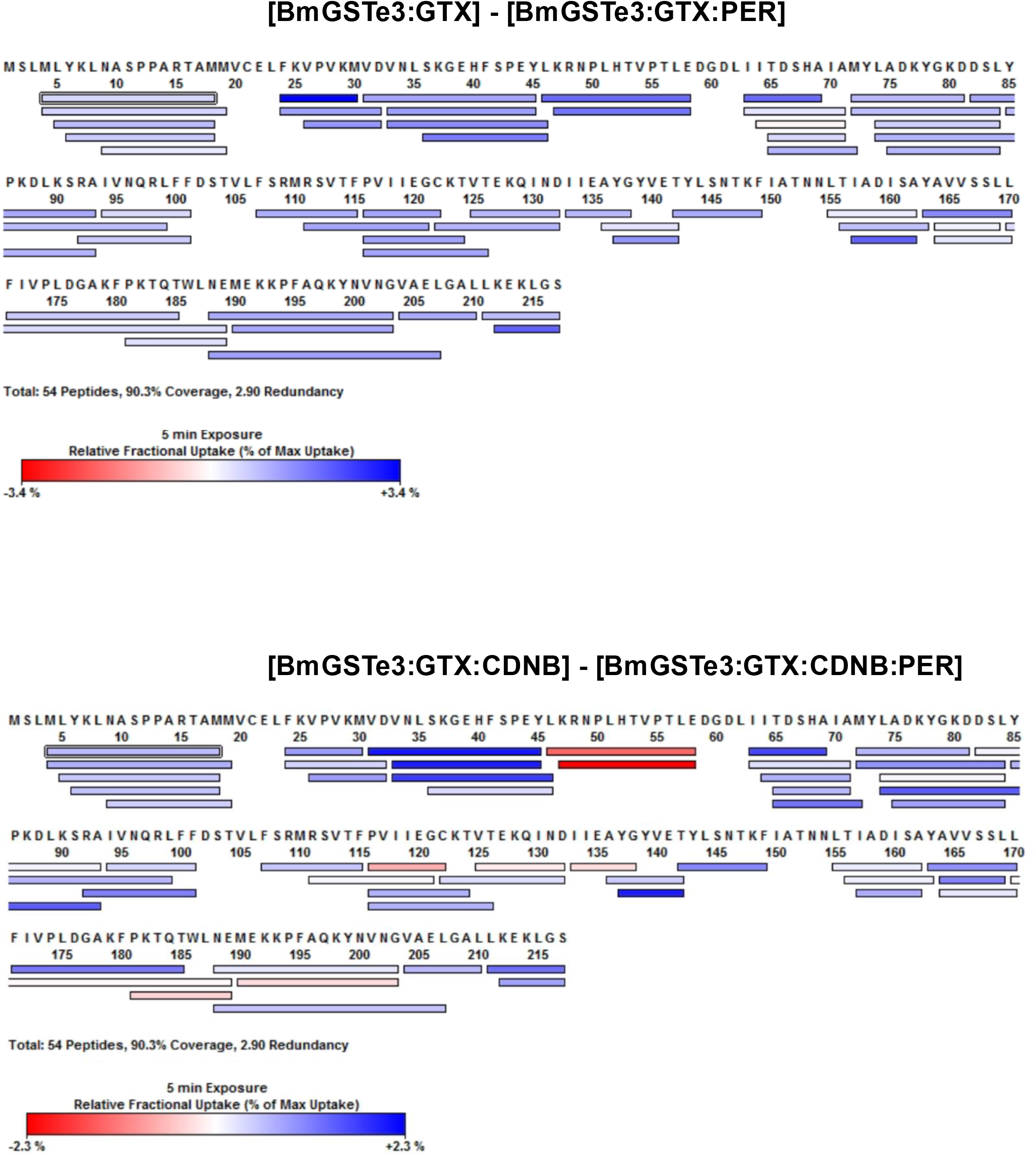
**Coverage maps of HDX-MS for individual states for BmGSTe3**

**Fig. S10.**
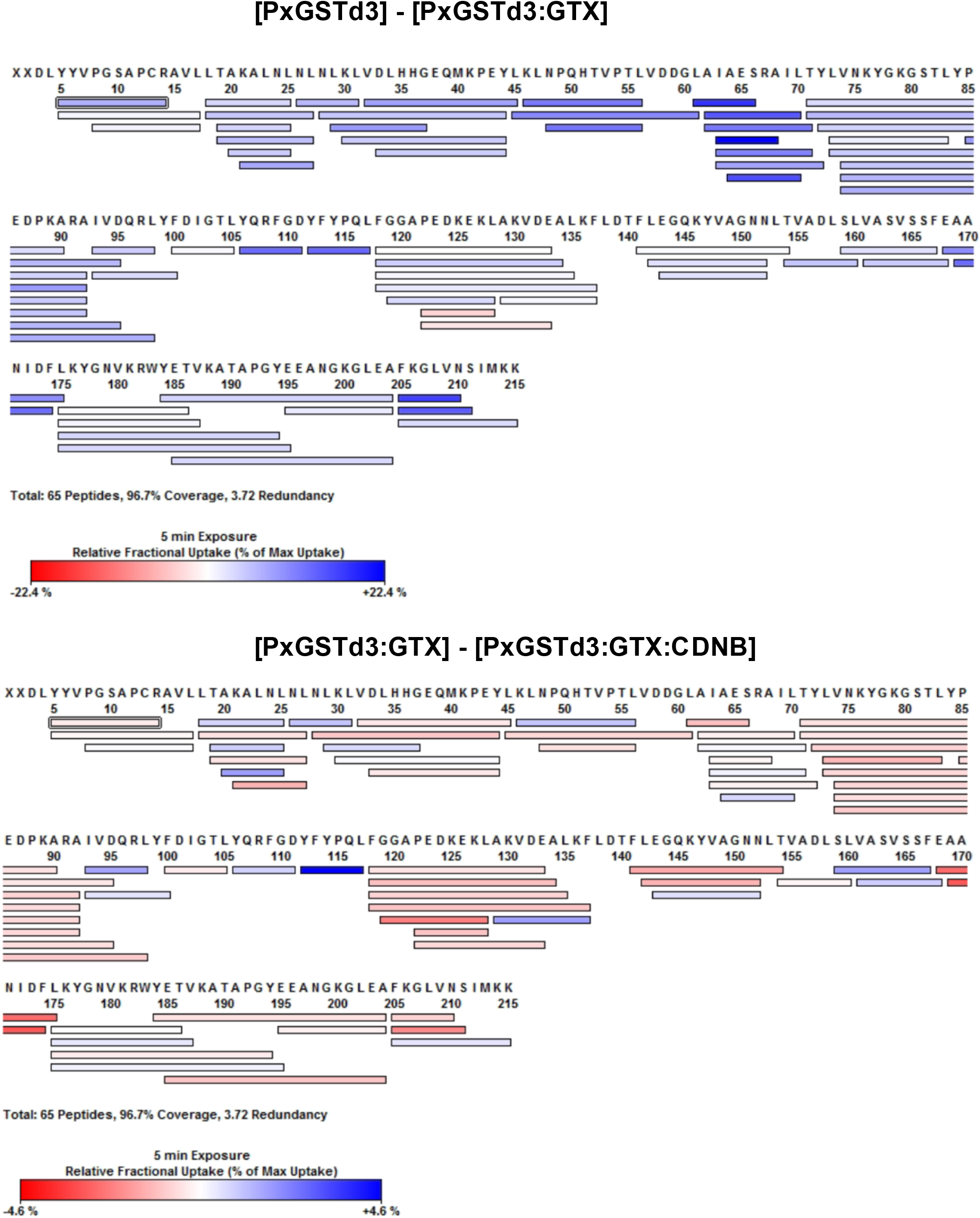
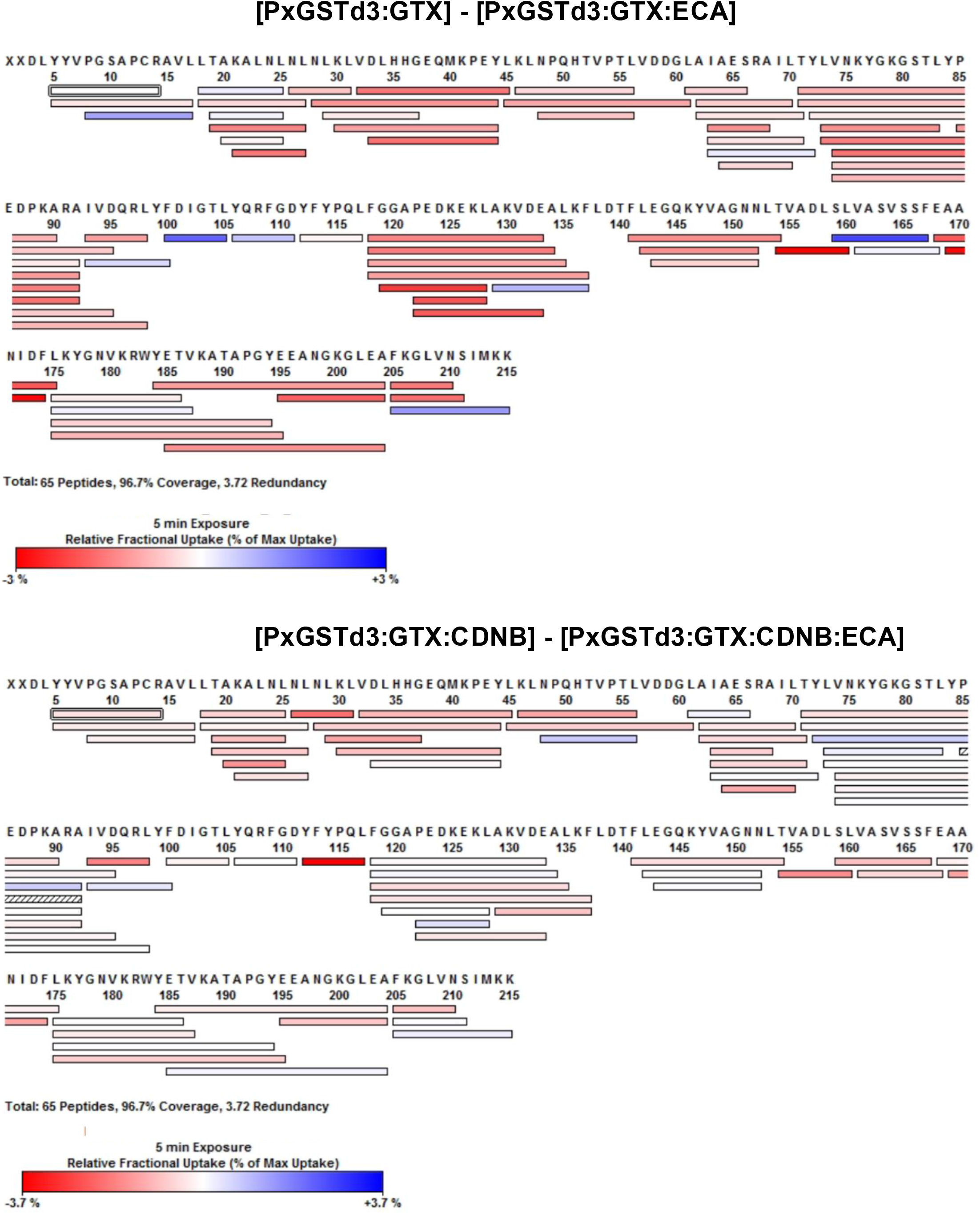
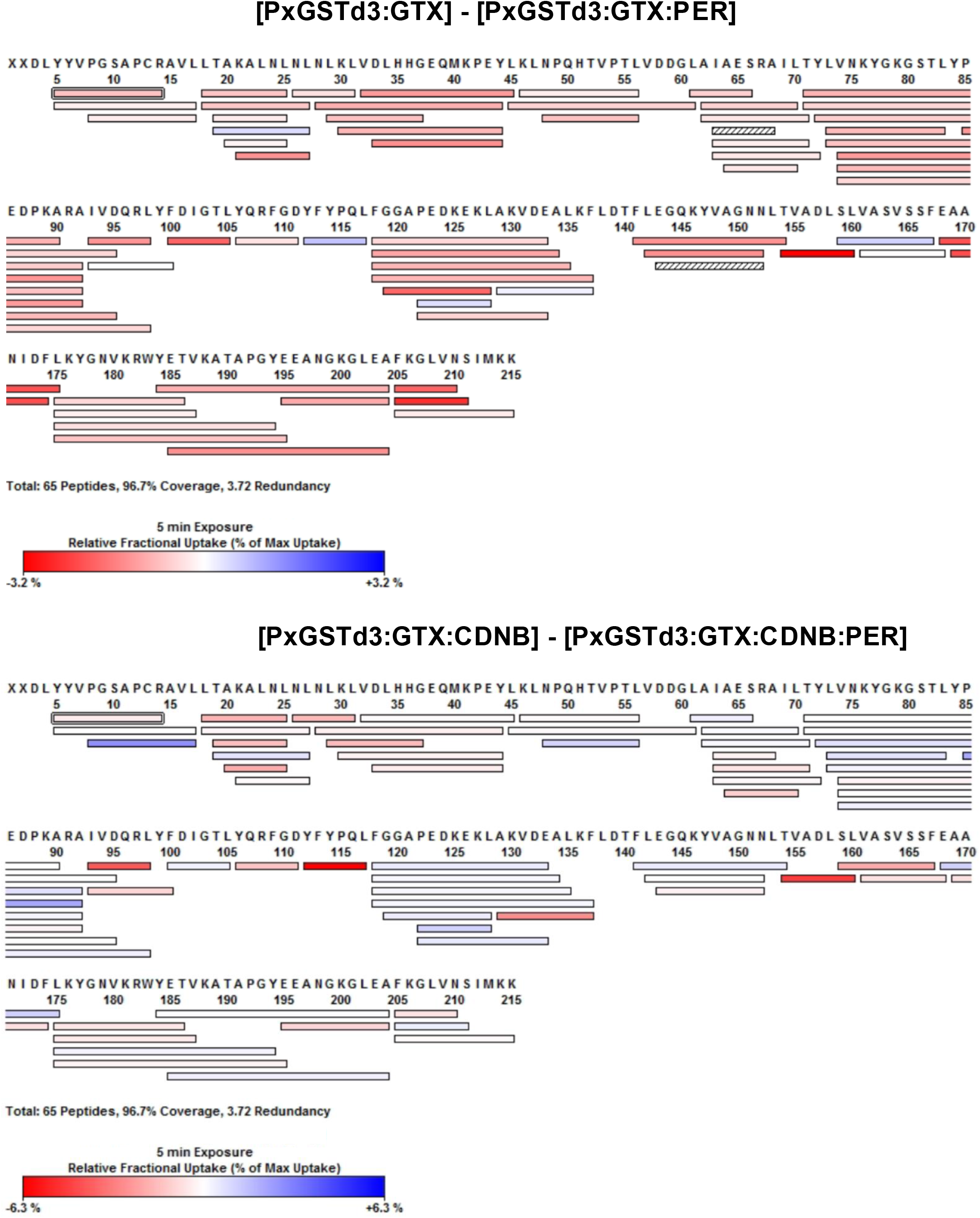
**Coverage maps of HDX-MS for individual states for PxGSTd3**

**Fig. S11.**
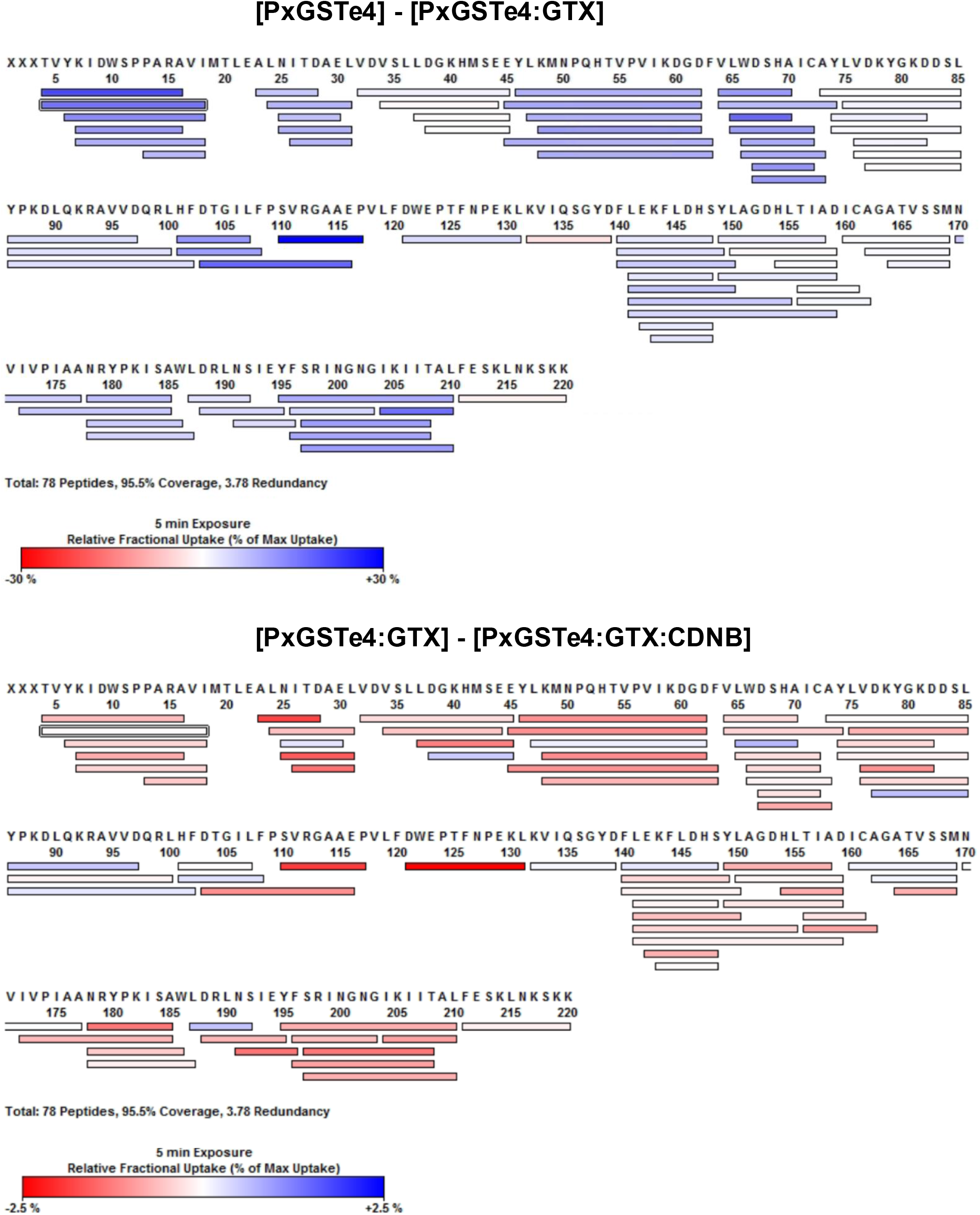
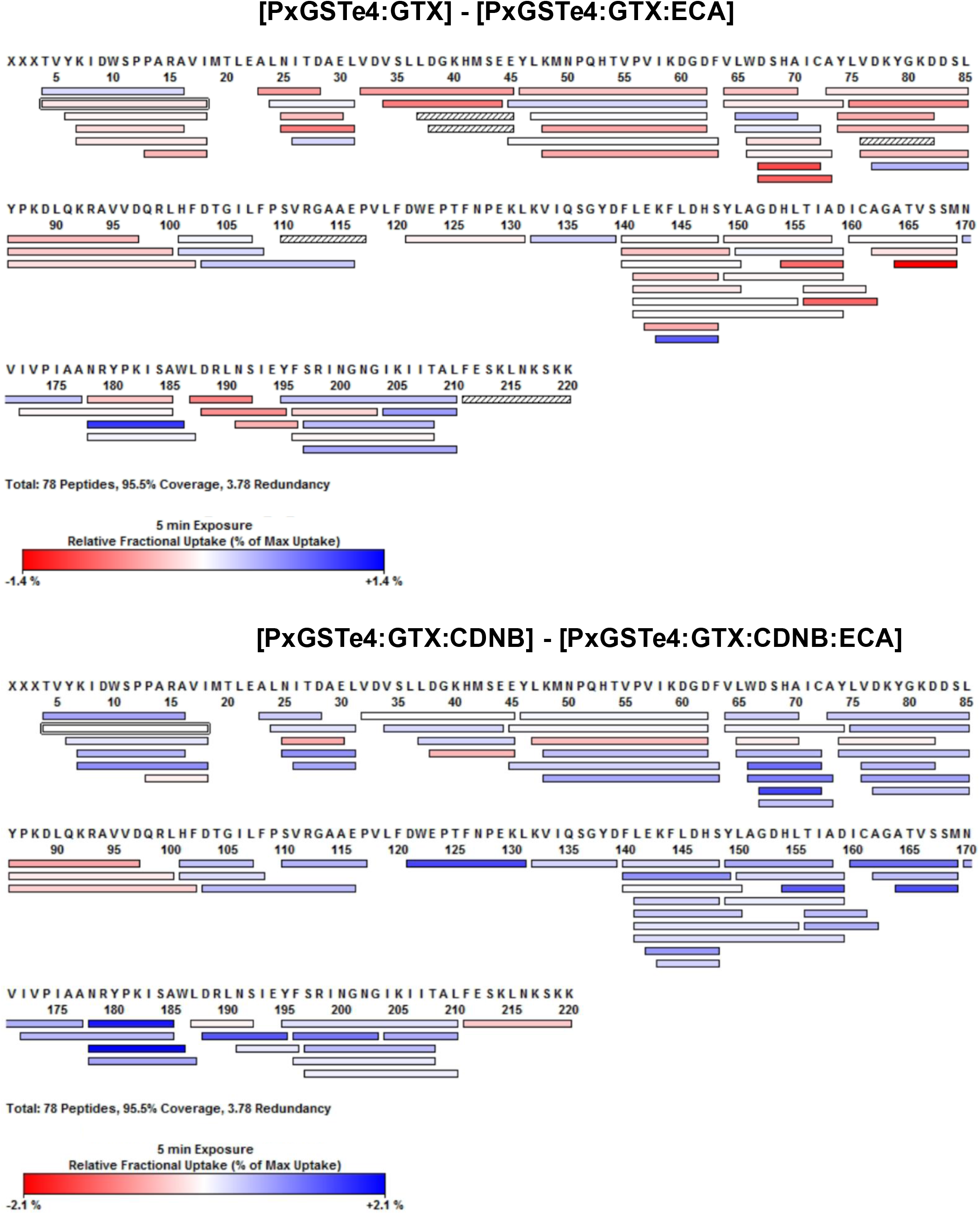
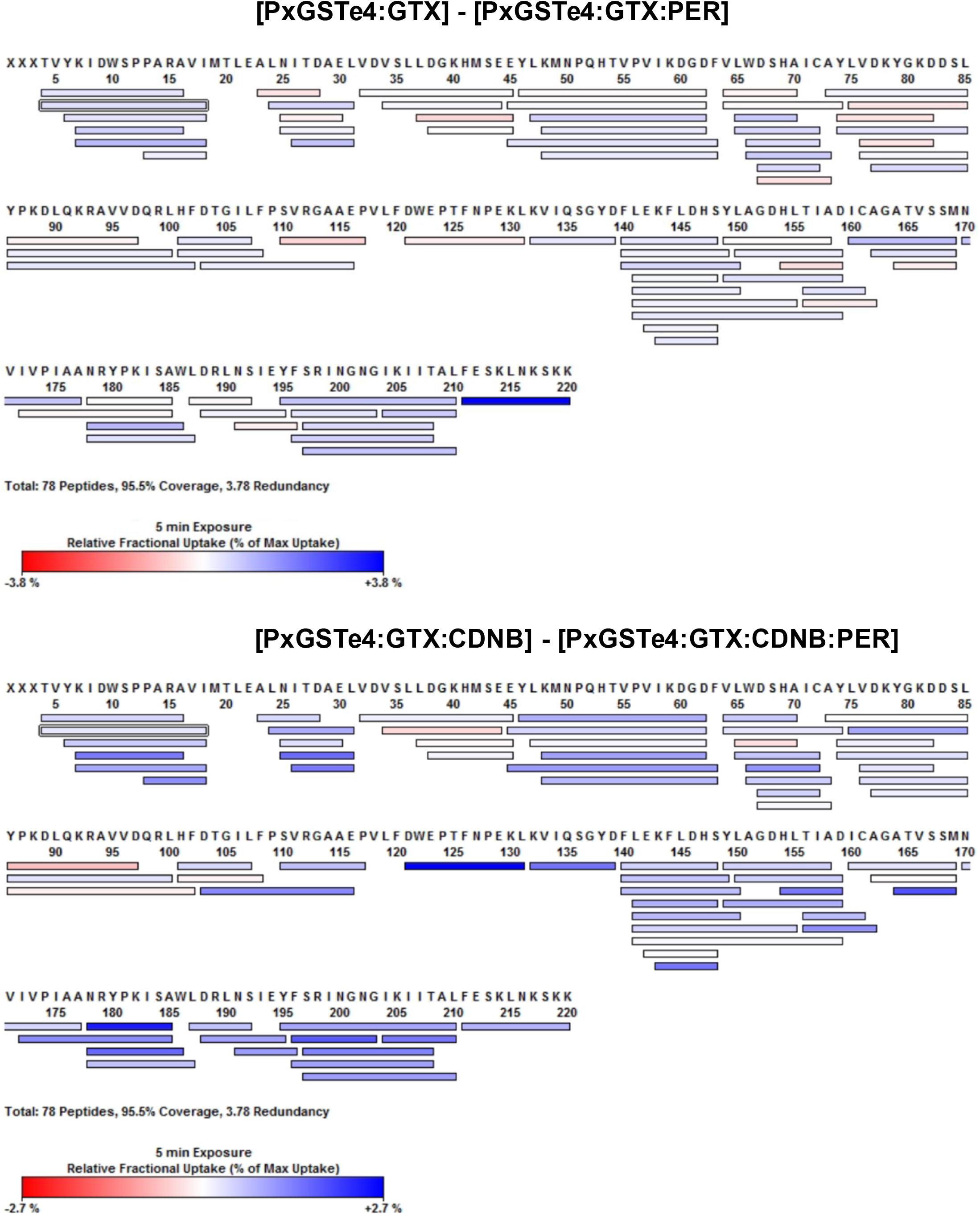
**Coverage maps of HDX-MS for individual states for PxGSTe4**

**Fig. S12.**
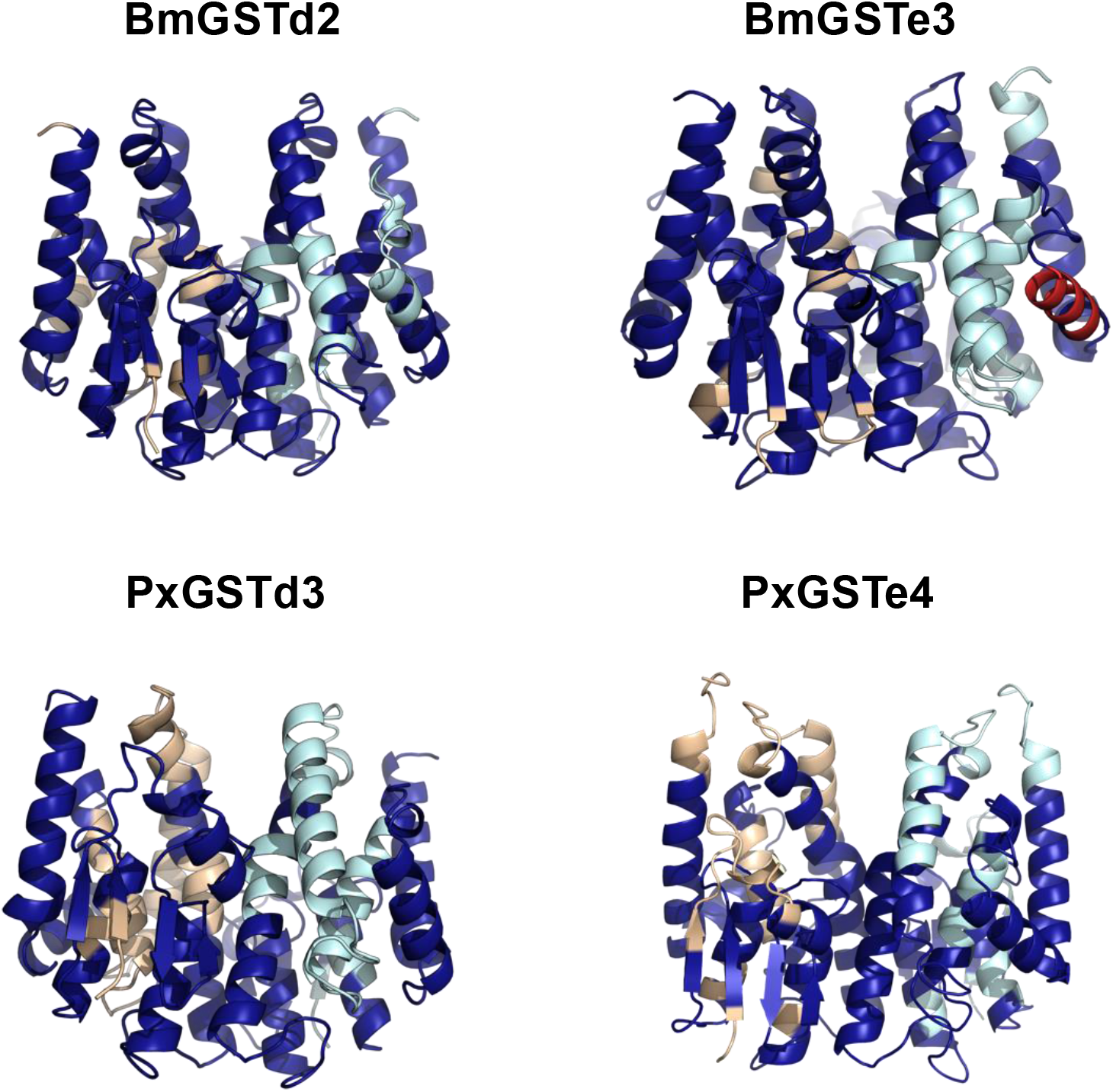
**HDX differences associated with binding of GTX to the selected apo-GSTs**

## SUPPLEMENTARY INFORMATION

**Supplementary Data File 1. Uptake differences in BmGSTd2 upon ligand binding**

**Supplementary Data File 2. Uptake differences in BmGSTe3 upon ligand binding**

**Supplementary Data File 3. Uptake differences in PxGSTd3 upon ligand binding**

**Supplementary Data File 4. Uptake differences in PxGSTe4 upon ligand binding**

## SUPPLEMENTARY TABLES

**Supplementary Table 1:** Structural alignments of AF3 models of target GSTs with consensus models and closest sequence homologues of known structure.

| Enzyme | Delta consensus<sup>b</sup> | Epsilon consensus<sup>b</sup> | 3F63 | 5FT3 | 7DB4 | 4E8E |
| --- | --- | --- | --- | --- | --- | --- |
| BmGSTd2<sup>a</sup> | 0.674 | 0.979 |  |  |  |  |
| LdGSTd3 | 0.809 | 1.148 | 0.759 |  |  |  |
| PxGSTd3 | 0.729 | 1.024 | 0.756 |  |  |  |
| ApGSTd1 | 0.797 | 1.035 | 0.727 |  |  |  |
| PxGSTe4 | 1.285 | 1.018 |  |  |  | 1.254 |
| BmGSTe3 | 1.228 | 0.955 |  |  | 1.001 |  |
| AaGSTe2 | 2.259 | 0.68 |  | 0.233 |  |  |

RMSD values are in units of Å.
<sup>a</sup>The crystal structure of BmGSTd2 has been solved (3VK9)
<sup>b</sup>AF3 models were generated from consensus sequences for the delta and epsilon enzymes in the sequence similarity network (Fig. 1a). Each target GST enzyme was aligned to both consensus models and also to its closest homologue of known structure (identified in the table by PDB ID).

**Supplementary Table 2:**
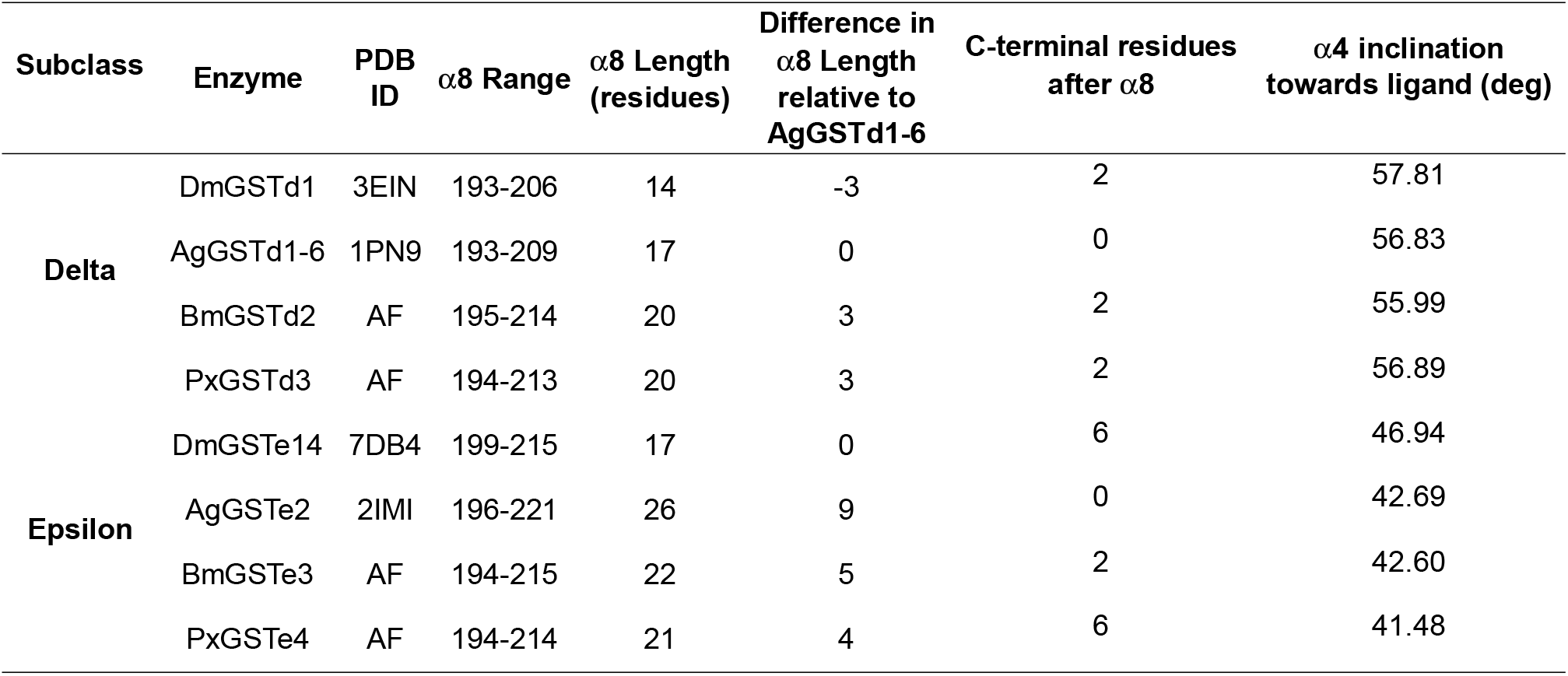
Differences in. α**-helix 8 (**α**8) length and** α**-helix 4 (**α**4) inclination**

| Subclass | Enzyme | PDB ID | α8 Range | α8 Length (residues) | Difference in α8 Length relative to AgGSTd1-6 | C-terminal residues after α8 | α4 inclination towards ligand (deg) |
| --- | --- | --- | --- | --- | --- | --- | --- |
| Delta | DmGSTd1 | 3EIN | 193-206 | 14 | -3 | 2 | 57.81 |
|  | AgGSTd1-6 | 1PN9 | 193-209 | 17 | 0 | 0 | 56.83 |
|  | BmGSTd2 | AF | 195-214 | 20 | 3 | 2 | 55.99 |
|  | PxGSTd3 | AF | 194-213 | 20 | 3 | 2 | 56.89 |
| Epsilon | DmGSTe14 | 7DB4 | 199-215 | 17 | 0 | 6 | 46.94 |
|  | AgGSTe2 | 2IMI | 196-221 | 26 | 9 | 0 | 42.69 |
|  | BmGSTe3 | AF | 194-215 | 22 | 5 | 2 | 42.60 |
|  | PxGSTe4 | AF | 194-214 | 21 | 4 | 6 | 41.48 |

**Supplementary Table 3:**
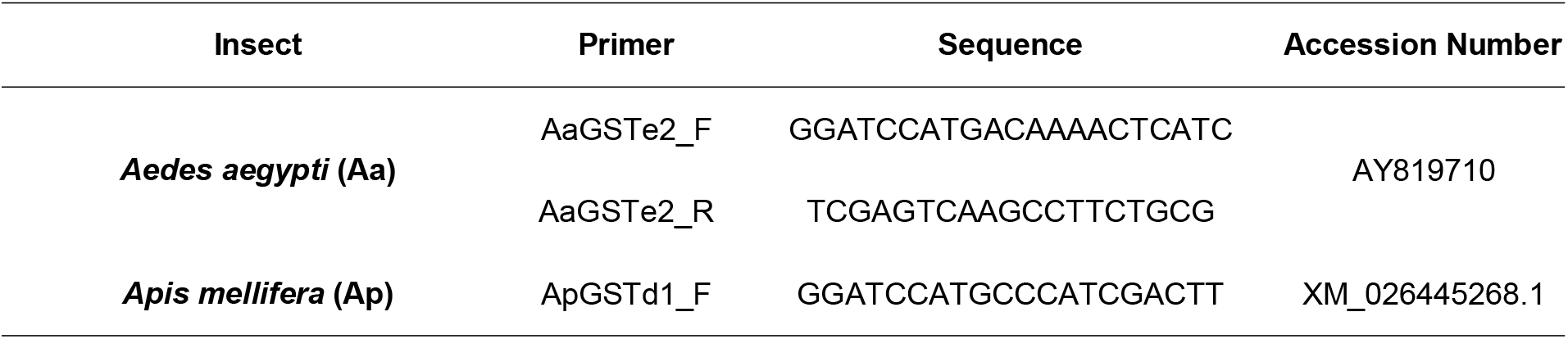
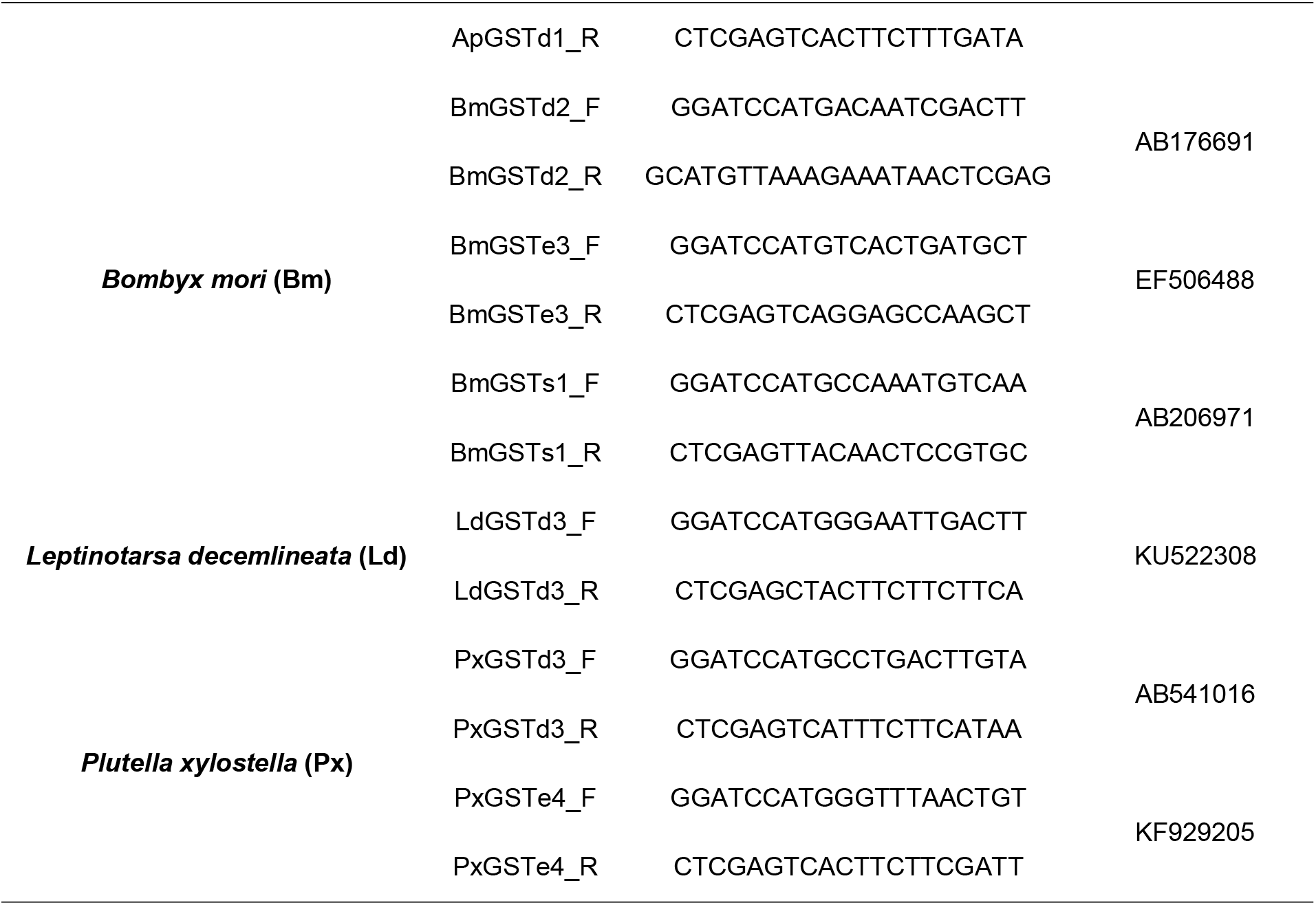
Primers used in this study.

## Notes

### Competing Interest Statement

The authors have declared no competing interest.

